# Engineering an in vitro spinal column: Manufacturing designs and emerging solutions for producing an axial mechanobiological system

**DOI:** 10.64898/2026.06.22.733686

**Authors:** Alexandra Iordachescu, Renush Vigneswaran, Aleksandar Atanasov, Liam M. Grover, Anthony D. Metcalfe, Aleksander Cendrowicz

## Abstract

The human spine is a complex, coordinated biomechanical system. Physiologically, its tissues are also highly interdependent in terms of function and viability. The interaction between mechanical stress and biological/biochemical activity over time constitutes a key driver of spinal degeneration.

Research to date providing mechanistic insights into this process has focused on individual components (vertebra and disc tissue analogues), in isolation or as basic functional units. However, many observations from individual units will not translate to whole spine behaviour. The intricate complexity of the spine requires novel experimental models (synthetic and biotic), which must consider the spine at an organ level and adopt an integrative approach that can capture the dynamics which govern its function.

Here, we report the development of a biomimetic spinal model prototype, amenable to cellular integration, which is miniaturised to the in vitro scale to provide a controlled environment and testbed for axial biological mechanics.

The research presented here encompasses more than a decade of systematic investigations during which the gradual emergence of key manufacturing innovations progressively enabled addressing an exceptionally complex bioengineering challenge – organotypic spine engineering.

The model comprises the full anatomical range of spinal vertebrae/bones (C_1_ to Sacrum & Coccyx), reproduced using bioceramic materials, assembled in sequence into a relevant columnar architecture and mechanically connected end-to-end by biochemically active interfaces. A range of assessments examining anatomical design, material behaviour and manufacturing processes is presented. The work explores concepts such as longitudinal mechanobiology and multi-segment coupling as well as manufacturing strategies using autonomous materials and instrumentation.

This prototype introduces for the first time columnar level behaviour and the ability to study time dependent adaptations. This model is important because it can support tissue maturation, evolving mechanical properties and adaptive behaviour and it represents an intermediate step between isolated skeletal tissue models and future organ-level spinal constructs.

## Introduction

The human spinal column (vertebral column) is a critical anatomical system, serving biomechanical and protective functions that are essential for survival. Forming the central axis of the skeleton, it is a multi-segment, multi-tissue structure which is evolutionarily adapted around the gravitational force, bipedal ambulation as well as dynamic mechanical load.

The spine is composed of osseous, ligamentous and fibrocartilaginous tissues connected in a manner which creates a flexible and curved structure (its characteristic sinusoid shape). This acts as a dynamic shock-absorbing system, enhances axial load distribution and maintains postural balance [1]. The S-shaped configuration reflects ontogenetic development as well as functional adaptations to movement patterns, with primary curves in the thoracic and sacral regions already existent at birth, while secondary cervical and lumbar lordotic curves emerge postnatally [2].

The spine comprises 33 bones during early development, of which 24 remain mobile in adulthood, as individual vertebrae separated by intervertebral discs (7 cervical, 12 thoracic and 5 lumbar vertebrae); and 9 bones/vertebrae become fused into two composite structures, the sacrum and coccyx (**Figure 1a**). The vertebral bones and intervertebral discs provide structure and support with internal loading forces and external mechanical forces, which increase in magnitude from the cranial to the caudal direction [3, 4]. This is translated anatomically through adaptations in the distribution and morphology of tissues, such as the increase in size from the cervical to the lumbar regions, or the architectures of vertebrae across the segments, reflecting local biomechanical demands [5]. Examples of regional adaptations include an increased mobility in the upper region, stability in the thoracic region and weight-bearing function in the lower spine, reflected in the vertebral bodies, spinous processes, laminae and pedicles shapes, orientations and sizes (**Figure 1a**).

**Figure 1:**
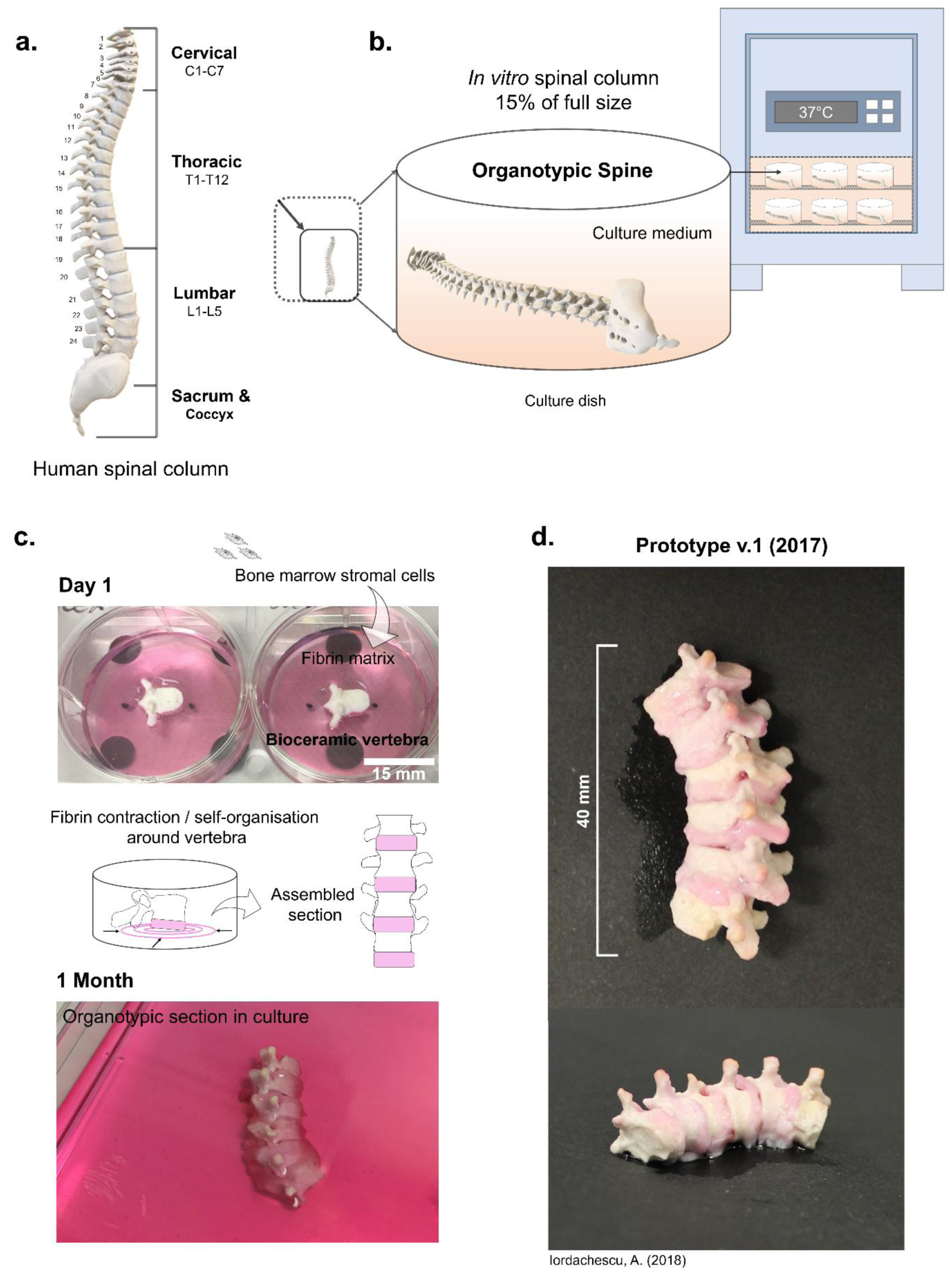
Conceptualisation and early development of an organotypic spinal model. **a** The spine represents the central axis of the human skeleton, which in adulthood comprises 24 mobile bones forming the cervical, thoracic and lumbar regions, and 9 bones fused into the sacrum and coccyx; connected by intervertebral discs in a flexible manner (a sinusoid shape). **b** Developing models of the spinal column as an integrated construct, at the in vitro scale (e.g. 15%) would be helpful for generating biotic models and studying developmental, adaptive and pathological processes. **c - d** A proof-of-concept biological prototype of a spinal column segment we developed between 2016-2017 to enable studying pathological spinal ossification. **c** Biomimetic ceramic vertebrae are used as anchoring points for self-assembling cell-seeded fibrin gels, which evolve to generate osteogenic interfaces. These are subsequently assembled in sequence (**d**) to generate a bioactive, columnar prototype (**Prototype v.1/2017**).

These highly interdependent tissues (vertebrae and discs) maintain each other’s spatial position, biochemistry and biomechanics, acting in a coordinated fashion, as a major anatomical functional complex. Whilst movement is limited between adjacent vertebra, acting as a unit, the spine is able to sustain a considerable range of motion [3]. Moreover, features such as shock-absorption depend on coupled deformation and collective activity across the different segments [4].

Physiologically, the spine also operates as a system. For example, the intervertebral discs are critically dependent on adjacent vertebral bodies and cartilaginous vertebral endplates for receiving vital nutrients. Their avascular structure means that they rely on the transfer of oxygen, glucose and metabolites from the vertebral circulation to support their matrices and remain viable [6].

The spine therefore functions as a continuous, highly-coordinated structure which exhibits a mechanophysiological behaviour that arises from the interaction of its subcomponents, particularly vertebral bones and discs. It is a kinetic system, where dysfunction within one of the tissues can disrupt the overall spinal biomechanics. For example, dysfunction within either the disc or vertebrae can initiate compensatory changes throughout the motion segments, altering load distribution throughout the whole column. As such, the spine is prone to a very high number of pathologies [4, 7], including degenerative disorders, traumatic injuries, congenital, neoplastic and inflammatory/metabolic disorders (**Figure 2**).

**Figure 2:**
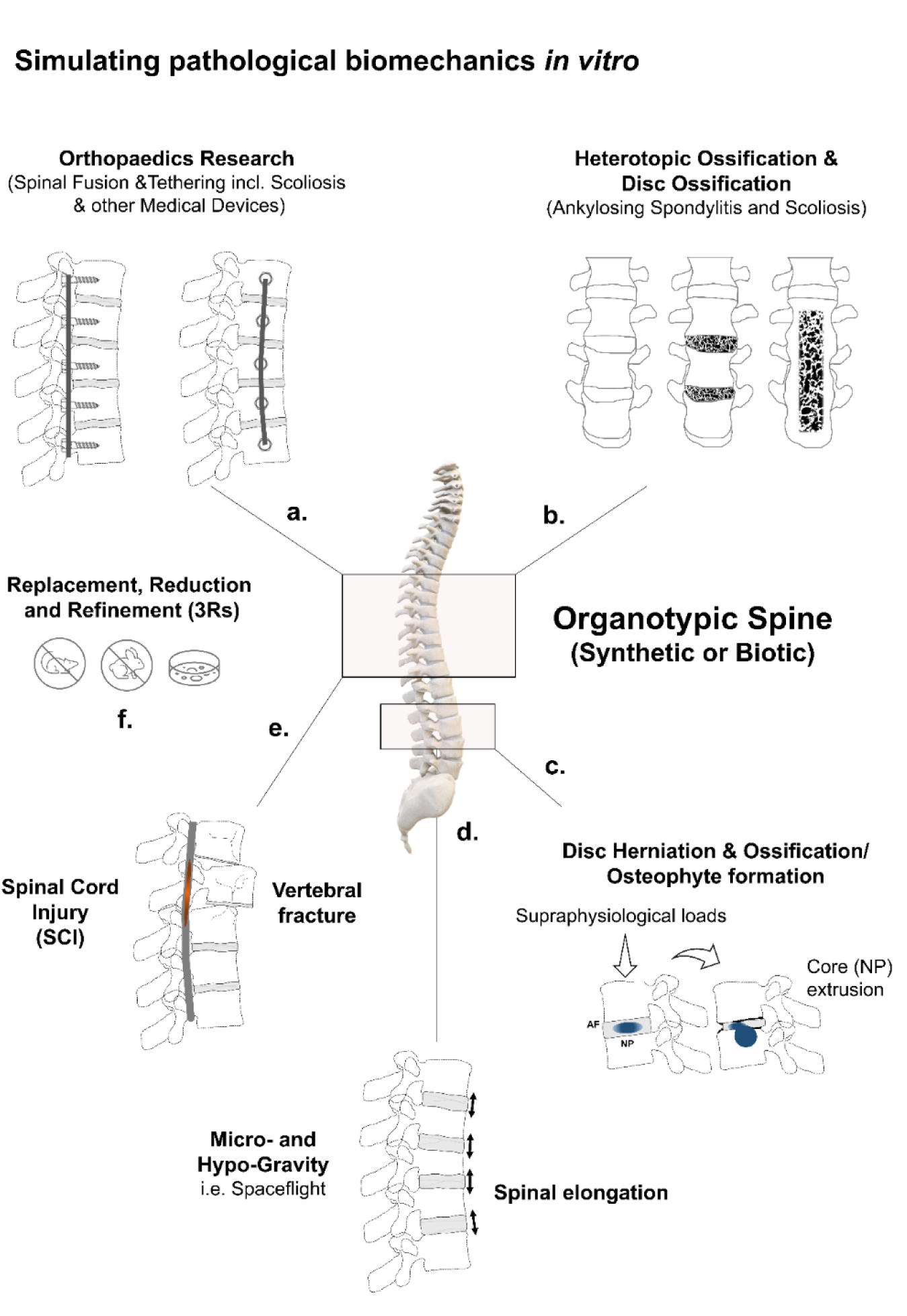
The increased demand for a tissue-engineered model of the spine. Bioengineering a multi-segment organotypic model of the spine (synthetic or biotic) can support research in orthopaedic/clinical materials (**a**), pathological biomechanics in spinal ossifications (**b**), disc degeneration and osteophyte formation (**c**), spinal elongation during axial unloading (**d**), vertebral fractures and associated spinal cord injury (**e**), and reducing the reliance on animal models (**f**).

Existing medical interventions for spinal degeneration are limited in their ability to restore the activity of the original tissue. Surgical interventions such as spinal fusion focus on spinal stabilisation of the damaged motion segments and restoring the local geometry/function [8], although they can inevitably affect surrounding segments, leading to maladaptive responses which accelerate degenerative changes at other spinal levels.

Studying these pathological processes and new interventions is particularly challenging because of the non-linear interactions between the tissues, the difficulty of simulating the physiological conditions in a laboratory setting and the reliance on donated patient tissue which can exhibit a high degree of variability. Many existing methodologies are focused on ex vivo mechanical testing of individual spinal components/functional units (single vertebra(e) & disc) [9], which do not necessarily reflect whole-spine behaviour. Furthermore, (patient) morphometric data reported across the extant literature shows large differences, reflecting characteristics which vary with demographic factors [5, 10]. Studying medical interventions or pathologies of the spine in other species (using animal models) is limited by significant anatomical, biomechanical and evolutionary differences, which are linked to different patterns of degeneration; as well as ethical considerations, as these models have a high severity index [11].

Computational model approaches such as finite element analysis (FEA) are the primary method that can be used to estimate the overall spinal behaviour under physiological loading conditions, pathology or surgical interventions [12, 13]. However, as with many modelling systems, their accuracy is dependent on the viability of the assumptions/parameters inputted as well as a simplification of certain tissue behaviours to make these computationally viable.

New experimental platforms to complement the in silico models would be highly desirable, as they could greatly enhance their predictive accuracy and help to gather novel evidence. Examples of emerging technologies include bioengineered and tissue engineered models, which at present mainly focus on producing intervertebral disc mimics [14–16]. However, no biological experimental platforms for the spine as a whole exist.

Developing bioactive experimental models of the spinal column as an integrated construct would be beneficial for biomedical, biomechanical and translational research. In particular, bioengineering models of the spine at the in vitro scale (containing living matter or otherwise) would enable generating organ-like-architectures, extracting quantitative biomechanical data, high-throughput experimentation and assessing biological responses/adaptation (in the case of cell models) (**Figure 1b**). This challenge is not simply additive (assembling engineered individual tissues together) but also systemic, requiring the identification of the parameters that are essential for the components to work together, which means the environmental settings, materials and instrumentation must be compatible. This manufacturing strategy (whole-organ engineering) would therefore enable the generation of a macroscopic architecture under a unified set of constraints.

As early as 2015-2016, we had begun considering how this multiscale modelling might be accomplished. In 2018, we reported a proof-of-concept biological prototype of a spinal column segment [17], composed of repeating bioceramic vertebrae connected in series by biologically-active interfaces, formed of self-assembling fibrin matrices seeded with bone-derived stem cells (mesenchymal stromal cells). We determined that a scaled-down range of 10-20% of the anatomical size was suitable for these applications. A 15% scale was identified as optimal, as it provided a balance between structural resolution and ease of manipulation. A further advantage was the ease of culture with respect to oxygenation and nutrient delivery/waste removal, which ensured the model could rely on passive transport to sustain cell viability, reducing the need for complex perfusion systems required in full-scale or explanted tissues (**Figure 1c-d**).

The aim of this exploratory work was to produce a biotic in vitro model of advanced pathological spinal ossification (as seen in conditions such as Scoliosis or Ankylosing Spondylitis). The research aimed to recreate typical phenotypes & morphologies encountered in such ossifications, where the intervertebral disc spaces are progressively to completely replaced by bone through abnormal osteogenesis mediated by cells including mesenchymal stromal cells [18]. This leads to longitudinal vertebral fusion into a single ossified element and to either flattening of the spinal curvatures or introduction of new deviations (**Figure 3c** provides examples of spinal deviations and malalignment).

**Figure 3:**
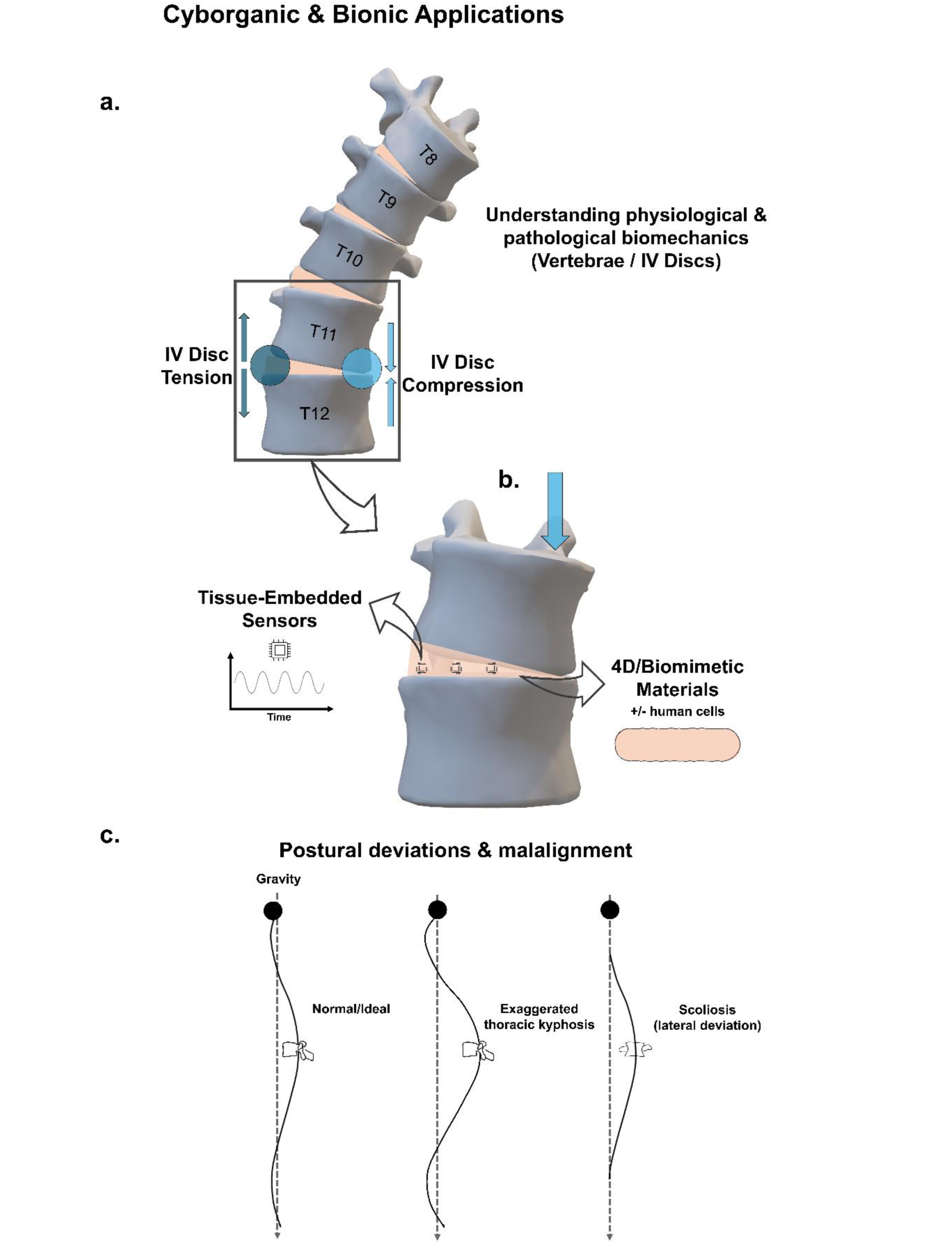
Engineering vertebral column analogues for emerging applications. Bioengineered spinal column models would be additionally useful as underlying substrates in bionic research models integrated with sensors, electronic or AI elements, which could provide real-time chemical/biomechanical measurements and analysis of physiological responses to mechanical stress (**a-b**). When integrated with morphing tissue mimics (with/without cells), they could help characterise maladaptive behaviour. **c** Postural adaptations and malalignment of the spine as a result of tissue ageing and deterioration (exaggerated thoracic kyphosis) or abnormal bioactivity leading to a pathological lateral deviation in Scoliosis.

The osteogenic interfaces connecting the vertebrae were developed based on a tissue engineered bone model which we had recently developed (now widely employed) [19–21] which could maintain viability during extended cultures (> 1 year) and evolve into mature bone. In this set-up, fibrin matrices, functionally analogous to the provisional matrix during tissue repair, when seeded with osteoprogenitor/stem cells, contract around biomimetic anchoring points, with anchorage determining emerging morphology and contractile organisation. In this case, the anchoring point was provided by individual, biomimetic mineral vertebrae, with bone marrow-derived mesenchymal stromal cells of murine origin integrated as the cellular component.

The spinal section reconstruction involved the assembly of the downsized (15%) anatomically shaped mineral vertebrae, developed by casting six replicas of the ninth thoracic vertebra (T_9_). The micro-vertebrae were reconstituted (as with previous models [19–21]) from a β-Tri Calcium Phosphate cement, a bioactive, osteoconductive ceramic used in orthopaedic procedures including spinal fusion; and which can evolve into the bone mineral hydroxyapatite in culture; as well as being a source of calcium and phosphate ions, facilitating the aberrant mineral deposition process within the interfaces. When placed individually inside 34.8 mm-diameter wells of culture microplates, each vertebra served as anchorage for fibrin matrices that were bio-actively generated in situ from the blood-derived constituents fibrinogen and thrombin. Because of the introduction of this single anchoring point, the cell-matrix constructs exhibited isotropic contraction towards the fixed, centrally positioned vertebra (**Figure 1c**), supported by traction forces generated by embedded cells, resulting in the formation of a compact, radially symmetric structure around the vertebral bodies.

The fibrin matrices - essentially biological adhesives - were also leveraged to provide an initial cohesion and gradual tissue integration with their respective and neighbouring vertebrae (throughout two months in culture). They are strongly adherent to bioceramics in culture, facilitating interlocking with the ceramic surface (which itself facilitates protein adsorption/adhesion); while simultaneously acting as temporary scaffolds for new tissue deposition (simulating the pathological state). Therefore, this design allowed not only the production of a continuous system but also one where the maturation/ biological adaptation of the system could be monitored (**Figure 1c**).

Once sufficiently contracted following an initial month in culture, a connected model was physically assembled using the 6 vertebrae and 5 connecting interfaces, which post-assembly measured approximately 40 mm in length (**Prototype v.1/2017** - **Figure 1d**). This organotypic construct was transferred to a 120 mm x 120 mm x 17 mm screening dish (144 cm^2^ base area) (**1c**) and maintained under standard, controlled culture conditions for an additional month to enable systematic observation (at 37°C, 5% CO_2_ atmosphere). During culture, the biological interfaces were fully submersed in culture medium for nutrition and media was changed regularly according to standard procedures. The work demonstrated that it was possible to maintain these constructs over extended cultures without contamination or media acidification, with medium pH remaining stable and matrix maturing as observed with normal bone constructs.

These early findings provided a foundation for the work reported here. In this research article, we describe the development of a higher-fidelity spinal model containing the complete vertebral set and axial structure/organisation, as well as disc surrogates which are amenable to cell incorporation. Our work systematically explores a range of advanced (bio)materials, manufacturing processes and physiological parameters which could enable whole-spinal organ engineering in the next decades.

This prototype introduces, for the first time, column-level behaviour and the spatial organisation of the spine (including axial architecture/regional structures), which is essential for multi-level mechanobiology, studying multi-joint mechanical coupling and long-term tissue evolution/maturation in a structured, continuous construct.

This prototype is significant as it constitutes an intermediate stage between extant, simplified prototypes (primarily tissue-engineered models of intervertebral discs) and more complex idealised systems that have yet to be realised; thereby providing a crucial development bridge towards increased model complexity by transitioning the field towards continuous, multi-segment spinal architecture.

## Results and Discussion

### The increased demand for a tissue-engineered model of the spine

Bioengineering a multi-segment organotypic model of the spine (synthetic or biotic) can support in vitro research on pathological biomechanics (**Figure 2**) such as investigating the development and refinement of medical interventions (e.g. spinal fusion and tethering) (**2a**); exploring the aetiology and pathogenesis of conditions such as Scoliosis and Ankylosing Spondylitis (**2b**), disc herniation and ossification/osteophyte formation (**2c**), spinal elongation during reduced mechanical loading (e.g. microgravity) (**2d**), traumatic vertebral damage or dislocation and the associated vertebral canal damage (including the encased spinal cord) (**2e**); as well as reducing the reliance on animal testing in orthopaedic research (**2f**). Engineered vertebral column analogues would be additionally useful as underlying models in emerging fields, such as cyborganic and bionic constructs integrated with sensors, electronic or AI elements. These hybrids could provide real-time chemical/biomechanical measurements and analysis of physiological responses to mechanical stress (**Figure 3a-b**). Potential applications include the integration of sensors/transducers/processing units with morphing/4D tissue mimics (with/without cells) (**3b**) to understand maladaptive behaviour encountered in postural deviations due to tissue ageing and deterioration (exaggerated thoracic kyphosis) or abnormal bioactivity leading to pathological malalignment in Scoliosis (**3c**).

### Producing bioactive vertebral and sacral-coccygeal mimics

The first phase of the spinal prototype development process involved manufacturing micro-vertebral templates with adequate geometries, surface topographies and resolutions that were appropriate for the scale of interest (15%, as determined previously) (**Figure 4**). These phantoms were used as intermediates in the manufacturing of final, biomimetic (bioceramic) vertebrae, which could recapitulate the rigid load bearing function of real vertebrae (**Figure 5a-c**).

**Figure 4:**
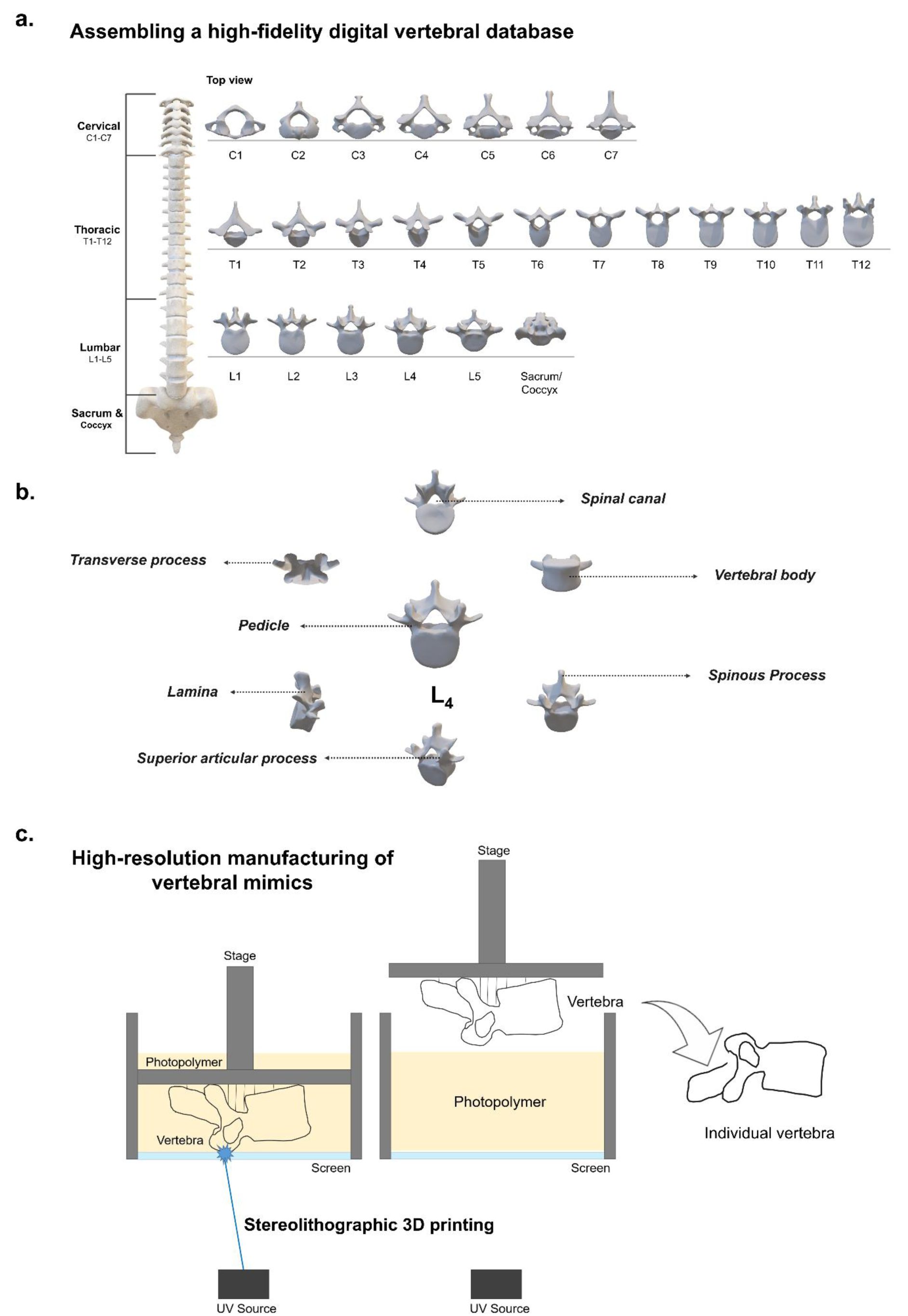
Producing high-fidelity vertebral and sacral-coccygeal phantoms. **a** A digital spinal database was assembled from an open-access collection, reconstructed from clinical 3D images and anatomical models. **b** These high-fidelity models contained the architectures specific to individual vertebrae, as illustrated using the fourth lumbar vertebra (L4). **c** A photochemical 3D printing technique was used to produce high resolution vertebral mimics at the reduced scale (15%).

**Figure 5:**
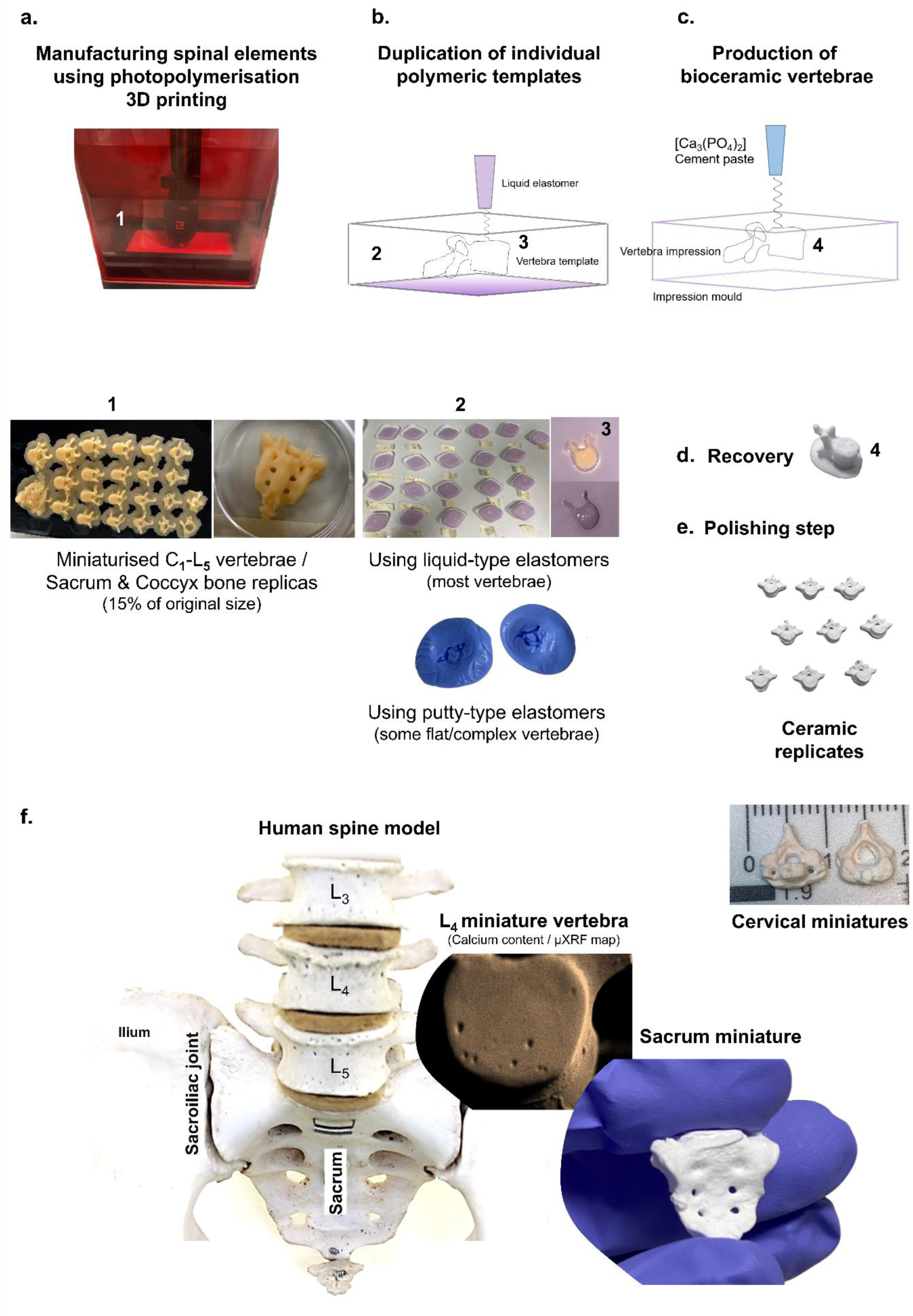
Translation of the biomimetic spinal components into bone-mimicking versions. a-c. The miniaturised anatomical phantoms of the complete range of spinal components were used as templates to generate final, bioceramic-based (β-TCP) vertebral and sacral-coccygeal versions using a combination of duplication and casting methods. These were subsequently recovered from their moulds (**d**) and where necessary, surface refinement was performed using mild abrasive treatment (**e**). The final structures showed strong morphological and topographical resemblance to corresponding anatomical structures in the human spine, as illustrated using the miniaturised L4 vertebra and Sacrum (**f**).

The first step involved assembling a digital spinal database, which contained true-scale 3D reconstructions of the complete range of human spinal bones - C_1_ - L_5_ vertebrae, sacrum and coccyx units (**Figure 4a**). These were sourced from a widely available open-access collection (see Methods section) containing idealised models developed from clinical 3D images and scanned models and measured against documented anatomical standards. These high-fidelity models contained the architectures specific to individual vertebrae (as demonstrated in **Figure 4b** for the fourth lumbar vertebra (L_4_)), including features such as the vertebral body, spinous process, superior articular processes, laminae, pedicles, transverse processes and the spinal canal. Some of these elements, such as the vertebral processes, are part of the spinal system which maintains alignment of the column, limits excessive movement and directs controlled motion. Therefore, replicating them in-vitro is important to simulate their function.

To manufacture the spinal structures at a high-resolution, a stereolithographic printing method (photopolymerisation 3D printing) was employed to 3D manufacture the complete set of spinal bones into high-fidelity polymeric duplicates of the digital counterparts (**Figure 4c**). This technology was able to reproduce the intricate structures, such as the spinous, transverse and articular processes at the reduced scale, something not achievable with additive fused deposition systems, both at that time (2019) and presently (2026) (**Figure 5a**). The technology uses the UV curing process of a photopolymer resin containing methacrylate monomers. The full set of vertebrae was fabricated in a single stage for consistency. The vertebral sets were additively printed in an inverted fashion when exposed to a 405 nm wavelength source, while emerging from a polymer (photoresin) bath (**Figure 4c**, **Figure 5a**).

The miniaturised anatomical replicates of the individual vertebrae, sacrum and coccyx bones were subsequently used as templates to generate biochemically relevant vertebrae using a combination of duplication and casting methods similarly employed in prosthodontics and dental reconstructions (**Figure 5a-c**). Individual vertebrae were immersed into either liquid-type commercial impression elastomers (most vertebrae) or putty-type elastomers suitable for flat and complex vertebrae (first cervical vertebrae, L_5_ and sacrum/coccyx) which were more difficult to duplicate/recover (**5b**). For example, the first two cervical vertebrae, the atlas (C_1_) and axis (C_2_) have atypical morphologies reflecting their function in providing greater mobility in the region.

The elastomers were allowed to cure for 24 hours (liquid elastomer) and 15 minutes (putty elastomer) to obtain an accurate three-dimensional impression of each vertebra/structure.

Owing to their comparatively low mass (in the range of tens to hundreds of milligrams vs. several grams), the polymeric vertebrae experience an upward buoyant force when immersed in the liquid elastomer, rising and remaining near the surface during the curing process. This convenient behaviour allows the vertebrae to be retrieved intact and without damaging the mould, as well as removing the need for additional processing steps. Furthermore, it creates an aperture which can serve as an access channel for the introduction of the liquid-state ceramic filling material.

Following curing, the vertebrae were carefully removed from the elastomeric moulds, leaving behind well-defined cavities mirroring their original geometry. These voids were subsequently filled with cement pastes generated, as reported previously [19, 21], through the reaction of the biomimetic ceramic powder (implant material) β-TCP (Ca_3_(PO_4_)_2_) with orthophosphoric acid (3.5 M H_3_PO_4_) (**5c**). This reaction between the mineral and protonating agent promotes a precipitation process and the conversion of the powder into a polycrystalline solid matrix. As identified in our previous work [19], the material is metastable in physiological conditions (i.e. cell culture media recapitulating the biochemistry of blood) and can transition to the prevalent bone mineral hydroxyapatite during culture. In our model, this material choice, formulation and chemical behaviour were important to produce a system that was sufficiently flexible to enable the study of biochemical maturation (and if used with cell substrates, biological maturation and tissue evolution).

Although the mineral-acid reaction is exothermic and cement setting occurs rapidly (over the course of minutes), the bioceramic vertebrae were allowed to set for a minimum of one hour to ensure uniform solidification and structural integrity, following which they were carefully extracted from their respective moulds (**5d**). Where necessary, surface refinement was performed using mild abrasive treatment to remove any surplus material and ensure batch uniformity across the replicates (**5e**). These final, bioceramic structures showed strong morphological and topographical resemblance to the corresponding anatomical structures in the human spine (**5f**). The average ceramic vertebral weight for the heaviest spinal vertebra, L_5_ (n=5) is 379.2 mg ± 6.68 mg, which represents approximately 2.12% of the average weight of a human lumbar vertebra (dry bone)[22].

A micro-X-Ray fluorescence spectroscopic elemental analysis (centred on Calcium and Phosphorus content) of ceramic replicates from batches duplicated/cast using the two types of elastomeric materials, revealed that the liquid-type elastomers, although characterised by longer setting times, yield a superior surface finish compared to the resin type, which cures rapidly (**Figure 6a-b**). However, resin models remain highly advantageous for accurately reproducing complex or more planar geometries and some intricate projections.

**Figure 6:**
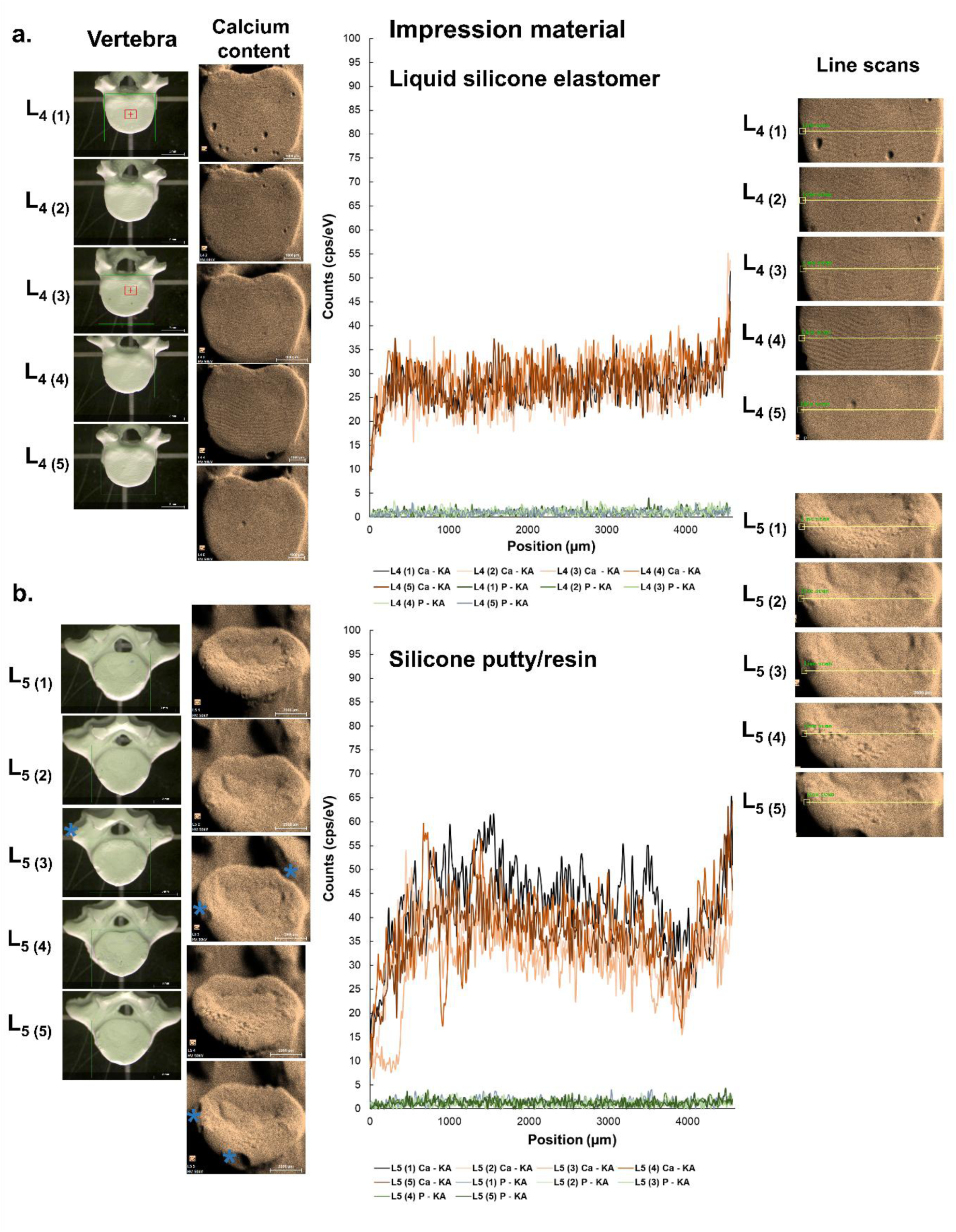
Characterising the manufacturing process using calcium phosphate mineral distribution in vertebral mimics. a-b. Micro X-Ray fluorescence elemental maps generated around Calcium content/distribution in samples; and spectra across line scans showing the distribution of Calcium and Phosphorus in five replicates from L4 and L5 ceramic vertebrae batches. **a** L4 ceramic replicates from batches duplicated/cast using a liquid-type silicone elastomer, used for most vertebrae. **b** Vertebrae with flat morphologies, such as L5, were duplicated/cast using a resin-type elastomer. Vertebrae produced using liquid-type elastomers display a superior surface finish compared to those produced using the resin-type. However, resin models are highly advantageous for accurately reproducing planar geometries. Blue stars represent minor abrasions incurred during processing rather than duplication.

### Whole-Column Engineering: Developing a multi-segment, coupled mechanobiological system

The next phase focused on reconstructing the human spine as an integrated system by building a controlled physical environment where axial bioactive structures could be studied as interacting, evolving networks rather than isolated components. The first stage consisted of the integration of the bioceramic vertebral components into a unified spinal architecture.

#### Modelling the spinal anatomy - establishing the columnar structure

The first step involved assembling the bioceramic micro-vertebrae C_1_ to L_5_ in sequence, in a columnar architecture, in a defined axis. Establishing the correct spinal conformation was essential for the production of this axial model. The human spine exhibits curvatures as a result of evolutionary optimisation, which absorb shock and dissipate forces during posture and locomotion. The regional curvatures of the vertebral column (cervical lordosis, thoracic kyphosis and lumbar lordosis) were mathematically assessed so that these geometric and functional features were accurately reflected in the model. This included reproducing the correct orientations at the sites of apophyseal joints across the regions, which can range from 45° in the cervical region, to 60° in the thoracic region and 90° in the lumbar region (**Figure 7a**).

**Figure 7:**
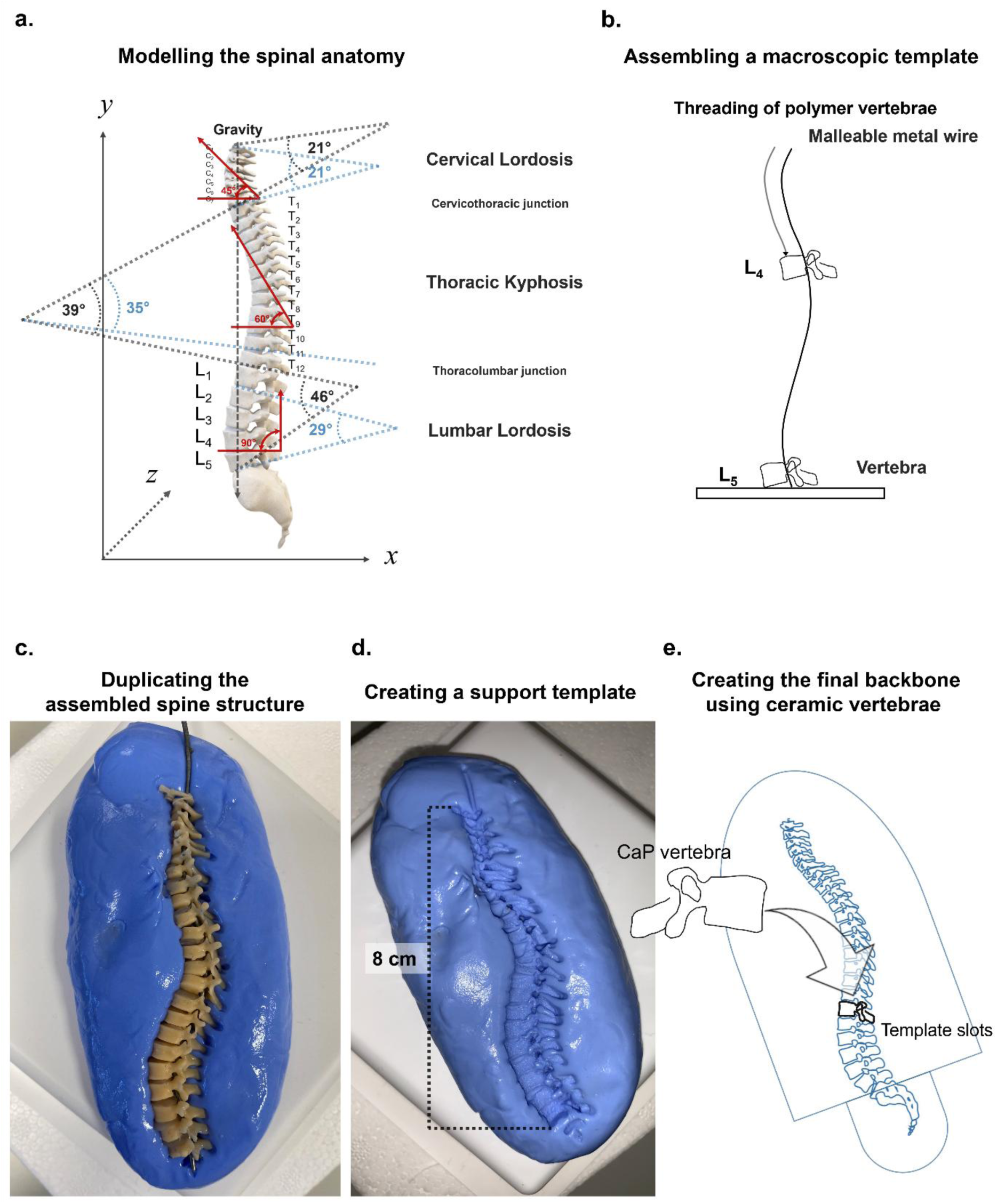
Whole-column engineering (1): Developing a vertebral backbone. **a** Modelling the spinal anatomy – the regional curvatures of the vertebral column were mathematically assessed in order to reproduce the correct spatial organisation and vertebral orientations. **b** A backbone was created according to these measurements using a malleable metallic wire. A macroscopic template was created by threading this structure through the spinal canal of the original polymeric vertebrae, which supported the vertebrae vertically, in the load bearing architecture. Pre-measured disc spaces were provisionally filled with orthodontic wax to prevent movement. **c** The overall structure was applied to a layer of resin-type elastomer to duplicate the assembled spine structure and enable the creation of a supporting template (**d**), which allowed inserting the ceramic vertebrae (**e**) and producing intervertebral discs in situ under more controlled conditions, negating some of the gravitational effects.

It is important to note that human spinal anatomy exhibits substantial variability across individuals, reflecting the complex interaction between genetic, lifestyle and environmental factors. Variations can include differences in vertebral number, asymmetries in vertebral facet joint orientations and deviations in the normal curvature and intervertebral disc heights. Because of this large natural variation, we developed a baseline model from which observations could be derived; and that could be easily modified according to individual research questions.

We measured and reproduced the relevant curvature angles in each region, based on ideal models reported in established medical educational literature [1], which helped to quantitatively define and reconstruct these anatomical features in the sagittal and coronal plane and at the reduced scale (**Figure 7a-b**).

For each spinal region of interest, the superior endplate of the uppermost vertebra and the inferior endplate of the lowermost vertebra defining the curve were identified. Angular quantification was performed by subdividing the spinal curve into smaller segments or selecting additional vertebral layers to account for the fact that physiological spinal curvatures do not completely conform to idealised arcs.

A backbone was created according to these measurements using a malleable metal wire (1 mm diameter), which was threaded through the spinal canal of the original polymer vertebrae and supported the set of vertebrae vertically, in the correct spatial organisation and load-bearing architecture, while the disc space heights were estimated and provisionally filled with orthodontic wax to prevent movement (**7b**).

#### Providing mechanical continuity - designing disc templates/interfaces

The subsequent step involved reproducing the role of intervertebral discs (IVDs) in the system by building connective interfaces which could enable assembly, spacing and mechanical continuity. A key consideration throughout the work was to build a system that was stable enough to maintain alignment, geometry and anatomical structure, but flexible enough to enable biochemical maturation and mechanical adaptation, as well as tissue remodelling if used with cells. This required the use of additional bioactive materials that contributed functionally to the system.

A wide range of biomaterials was considered, which could function as tissue analogues to produce a system that was mechanically connected end-to-end (C_1_ - Sacrum & Coccyx). These included natural biomimetic polymers (e.g. polysaccharides and protein/ECM-derived matrices), synthetic and composite materials, aiming to find a balance between favourable biological properties, mechanical performance and adaptive properties/autonomous behaviour, as well as ease of manufacture. Additional considerations included cellular compatibility; their microstructure and degree of structural modifiability to simulate the anatomical organisation of an IVD into its two components (outer, *annulus fibrosus* and inner, *nucleus pulposus*); biomanufacturing feasibility; as well as the extent of their established role as matrices in tissue engineering.

Based on these factors, a formulation for a multi-phase disc interface was developed composed of the bio-derived, water-soluble polysaccharide gellan gum (low acyl, in a fluid gel form) encapsulating the thermoresponsive biopolymer gelatine. Both material formulations generated polymers with useful, autonomous life-cycle control. These interfaces were designed to replicate the mechanical and organisational roles of an IVD, including providing a compliant load transfer zone simulating the IVD role in absorbing and distributing the spinal load.

Gellan gum (C_24_H_38_O_20_) is a biosynthesised polysaccharide which is increasingly significant in several industrial applications due to its gel forming abilities with tuneable rheological and biomechanical properties, meaning it can be adapted to generate bespoke extracellular matrices for tissue engineering and regenerative medicine. A linear, anionic polysaccharide, it is composed of repeating tetrasaccharide units containing the sugar residues β-D-glucose, β-D-glucuronic acid and α-L-rhamnose in the [Glc-GlcA-Glc-Rha]_n_ conformation [23]. Here, this polymer was exploited for its pseudoplastic properties, which made it possible to encapsulate another material phase, as well as being advantageous for 4D biomanufacture through its stimuli-responsive, self-healing abilities. When prepared as a non-Newtonian fluid gel (1.5% w/v concentration in dH_2_O) through a shear-cooling production process, it could be injected into the narrow disc spaces to occupy and adapt to those geometries more successfully compared to typical gels.

Xenogeneic gelatine (partially hydrolysed collagen; of suine origin) at a concentration of 3.7% w/v was used as a simple physical analogue for the *nucleus pulposus*, to mimic the hydrated, protein rich composition related to its mechanical properties, which depend on large molecules that bind water and form flexible networks (**Figure 12f**). Gelatine is a polymeric substance containing the amino-acids glycine, proline and hydroxyproline [24–26] which are also abundant in collagens, including types II and XI, present in the *nucleus pulposus* (as a fibrous meshwork supporting a proteoglycan gel including aggrecan; or the *nucleus pulposus* - *annulus fibrosus* interface). These amino acids are also building blocks for aggrecan, which is rich in glycine and proline [27]. Structurally, gelatine gels form thermoreversible protein networks through the partial renaturation of collagen polypeptides, which can help model the *nucleus pulposus* behaviour during culture at 37°C (the core body temperature) to maintain a semi-fluid morphology. Gelatine is also a widely used material for biomedical applications [28, 29], due to a significant genetic and proteomic homology between many species and human collagens.

Therefore, gelatine was selected for constructing the *nucleus pulposus* core due to its physiological relevance, to mimic the viscoelastic behaviour, water retaining properties and protein-rich content of this inner structure, while the encapsulating *annulus fibrosus* was produced using gellan, due to its spatiotemporal behaviour, biocompatibility and mechanically controllable properties. These aspects are further elaborated in the next sections.

#### Vertical integration and accounting for gravitational effects

Gravity is a particularly important biophysical consideration both in manufacturing this system (during vertical construction) and post-assembly, as it can influence the organisation (and subsequent evolution) of the constructs. This includes mechanotransduction arising from gravity-induced effects such as compressive strain due to weight forces as well as hydrostatic pressure within the fluid-filled regions of the construct (in discs).

Vertical integration of the discs into the spinal model was initially trialled in a sequential manner, starting from L_5_. However, this proved challenging due to gravitational effects, given the low stiffness of the disc templates, which were susceptible to deformation under the vertebral weight before sufficient setting had taken place to stabilise their geometry. Therefore, a stepwise vertical assembly was not efficient and increased the risk of misalignment. A bespoke methodology was created, where the vertebrae could be first positioned into place, allowing for the discs to be introduced and set under more controlled conditions, in one stage, thereby reducing the likelihood of damage and ensuring uniformity across disc constructs.

To achieve this, the spine as a unit was imprinted into a layer of impression resin to duplicate the assembled spine structure (**7c**), creating a whole-spine support model containing accurate template spaces for bioceramic vertebral insertion and enabling subsequent disc generation (**7d-e**).

The bioceramic vertebrae were subsequently inserted into the template spaces, by fitting firstly the bodies and subsequently aligning the spinous processes within the corresponding location, creating a final backbone. The spaces between the vertebrae were therefore preserved to allow insertion of disc matrices. The range of intervertebral disc heights was estimated based on adjacent micro-vertebral heights (derived from the polymer vertebrae). These are presented in **Figure 8a**, with the intervertebral disc heights accounting for 25% of the mobile spine length and approximately 20% of the total vertebral column, consistent with normal anatomical structure [1, 30].

**Figure 8:**
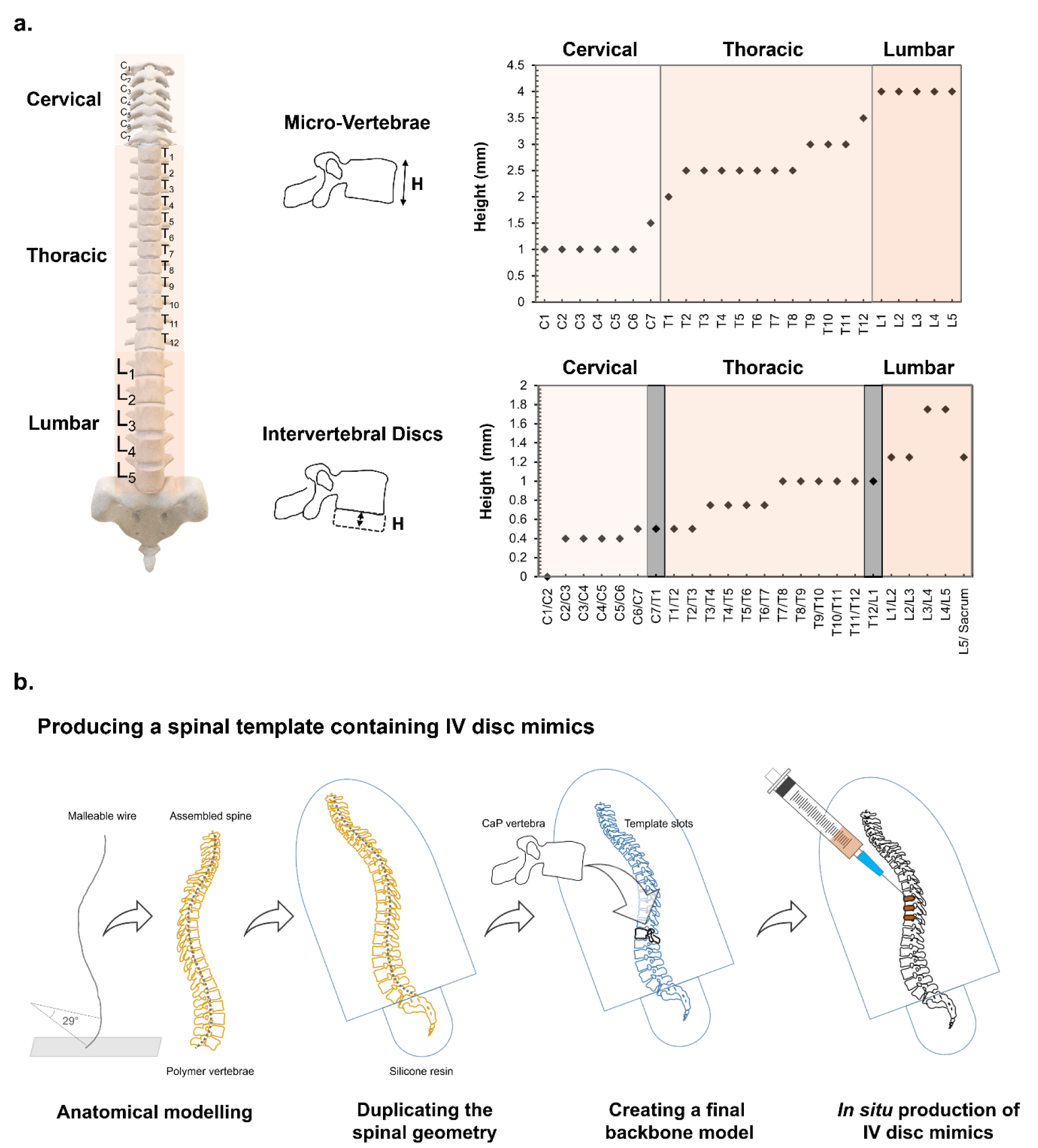
Producing the disc interfaces in situ. The range of intervertebral disc heights was estimated based on adjacent micro-vertebral heights (**a**). The surrogates for the intervertebral discs (engineered biopolymers) were subsequently injected in situ, into the spaces between adjacent vertebrae and between the final mobile vertebra (L5) and sacrum, to generate an integrated, mechanically-connected spinal model (**b**).

#### Producing the disc interfaces *in situ*

The surrogates for the intervertebral discs were subsequently injected *in situ*, into the spaces between adjacent vertebrae, and between the final lumbar vertebra (L_5_) and the sacrum/coccyx bone mimic, which had been previously sized to the anatomically correct heights. A formulation of the polysaccharide gellan (low acyl,1.5% w/v concentration in in dH_2_O) was produced as a non-Newtonian, fluid gel version through shear-cooling and used as the scaffolding material. The material was labelled with a trace of Alizarin Red S pigment to enhance visual contrast, due to the high optical transparency of gellan gels. Based on the morphometric measurements, it was calculated, by considering the anatomical disc compartments as concentric cylinders with volumes V_IVD/NP/AF_=πr^2^h, that up to 0.05 ml of polymer would be required for each disc in the lumbar region; 0.03-0.04 ml in the thoracic region (mid-upper thoracic and mid-lower thoracic, respectively); and 0.2 ml in the cervical region (**Figure 8b**, **Figure 9a**). Following application into the IVD spaces, the gellan structures were allowed to rest for approximately 30 minutes, for the gel networks to reform/reorganise from a flowing dispersion during injection/shearing.

**Figure 9:**
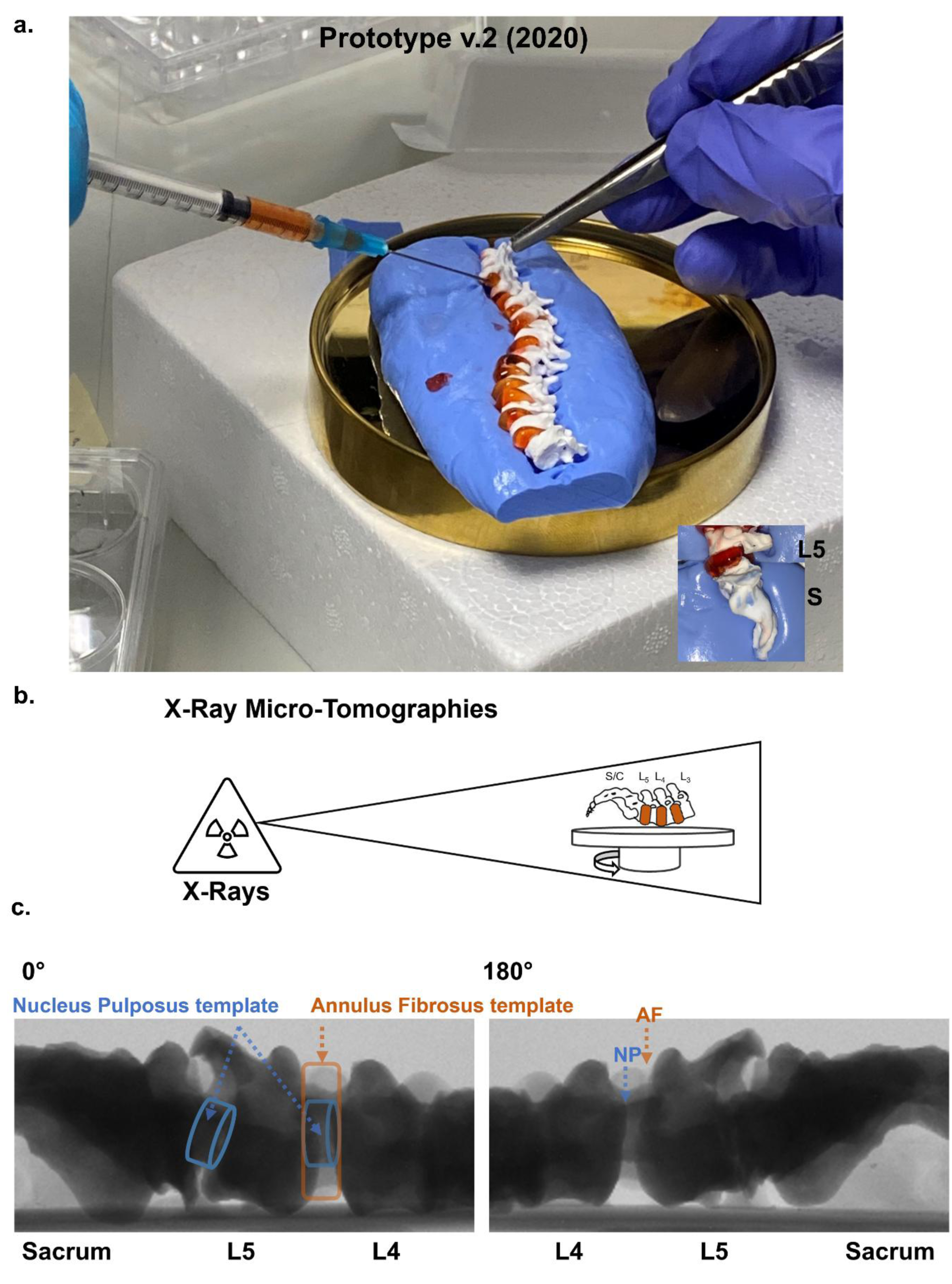
Whole column engineering (2): Developing a multi-segment, coupled mechanobiological prototype. **a** The final prototype, (**Prototype v.2/2020**) developed between 2019-2020, consisted of a vertebral backbone, mechanically connected end-to-end by multi-phase biomimetic discs, developed by engineering the rheological/mechanical properties of the bioderived polymers gellan gum and gelatine. The intervertebral disc heights (**a**) accounted for approximately 20% of the total vertebral column, in line with normal anatomical structure. To assess the viability of the production process, the lumbar-sacral region, containing the largest discs, was detached from the rest of the spine and subjected to X-Ray analysis using a micro-computed tomography system (**b**). The X-Ray images in **c**, focused on the L4-L5 interface, acquired at different rotation angles, show the successful compartmentalisation of the internal nucleus, consisting of gelatine (simulating the *nucleus pulposus*) within the suspending gellan shell (used as a surrogate for the *annulus fibrosus*).

To simulate the structure and bioactivity of the jelly-like inner core of intervertebral discs, the *nucleus pulposus* was modelled using the biopolymer gelatine (3.7% w/v in dH_2_O), a useful building block due to its ease of adaptability and functionalisation; the presence of adhesion motifs for cells, and biocompatibility. These hydrogels can be further engineered to match the viscoelastic properties of the *nucleus pulposus* and in a biological scenario, can support the cell production of relevant proteoglycans and collagen species. Their injectability, as well as *in situ* gelation, were their most practically important advantages in this system. Furthermore, similarly to fibrin, used in the first prototype (**v.1/2017**), their degradability can be tuned to match new tissue formation.

Gelatine cores were injected using a 1 millilitre hypodermic syringe connected to a 23-gauge needle, into the central portion of the gellan *annulus fibrosus* templates, loading a volume equivalent to 40% of the original V_AF_ and occupying approximately 30% of the total disc volume (V_IVD_), in line with a typical healthy disc [31] (**Figure 12f**). The supporting *annulus fibrosus* templates were allowed to self-organise/self-heal around the encapsulated structures.

These intervertebral disc analogues were therefore integrated into the columnar structure as functional surrogate interfaces between the vertebral segments. They were designed to replicate key mechanical and organisational roles of a disc, including the load distribution element, acting as hydrated, compliant materials.

The final, engineered spine (**Prototype v.2/2020)** measured approximately 10 cm in length, with the mobile (presacral) spine measuring 8 cm (**Figure 9a**, **Figure 7d**). This represents approximately 15% of an average 70 cm (27.5 inches) human spine, often used as a standard for human morphological data [1].

To assess the viability of the assembly process, the lumbar-sacral region containing the L_1_-Sacrum & Coccyx mimics and disc interfaces, was detached from the rest of the spine and subjected to X-Ray analysis (**9b-c**) using a micro-computed tomography system, to visualise the internal structures based on their radiodensity. This regional scan, as opposed to a whole-spine scan was chosen in order to achieve micron-scale imaging resolution at the tissue scale (large field of view). This section of the spine was chosen due to its essential role in weight-bearing. This assessment provided insight into the structural behaviour of this multi-component segment, including the containment of the nucleus within the suspending disc post-assembly, while supporting the vertebral/internal weight unassisted (through cohesive and adhesive forces and internal strength).

The X-Ray images in **Figure 9c**, acquired at rotation angles of 0° and 180°, show the successful compartmentalisation of the internal nucleus within the *annulus fibrosus* template, as seen in this example between adjacent vertebrae L_4_ and L_5_. Throughout the handling and imaging process, the spinal segment maintained cohesive stability and the components remained assembled without any displacement or separation.

Overall, this set of experiments confirmed the feasibility of producing a load-bearing architecture in a biochemically active, axial system. This is conceptually important because it offers an extension to functional spinal unit models (single/two vertebra(e) + connecting disc) by introducing multi-level coupling and interactions across a chain of vertebrae.

This baseline model can be tuned with additional bioactive materials, cellular content and structural reinforcement to increase anatomical complexity and observe adaptive behaviours in a mechanically/gravity loaded system.

Furthermore, the mechanical properties of the *annulus fibrosus* gellan template can be tailored due to its polyelectrolytic nature, through ionic crosslinking in the presence of a Ca^2+^ rich solution, to transition the fluid-like state into a structured, bulk gel network, thus optimising stiffness and its ability to act as a scaffold for cells (**Figures 10 and 12**).

**Figure 10:**
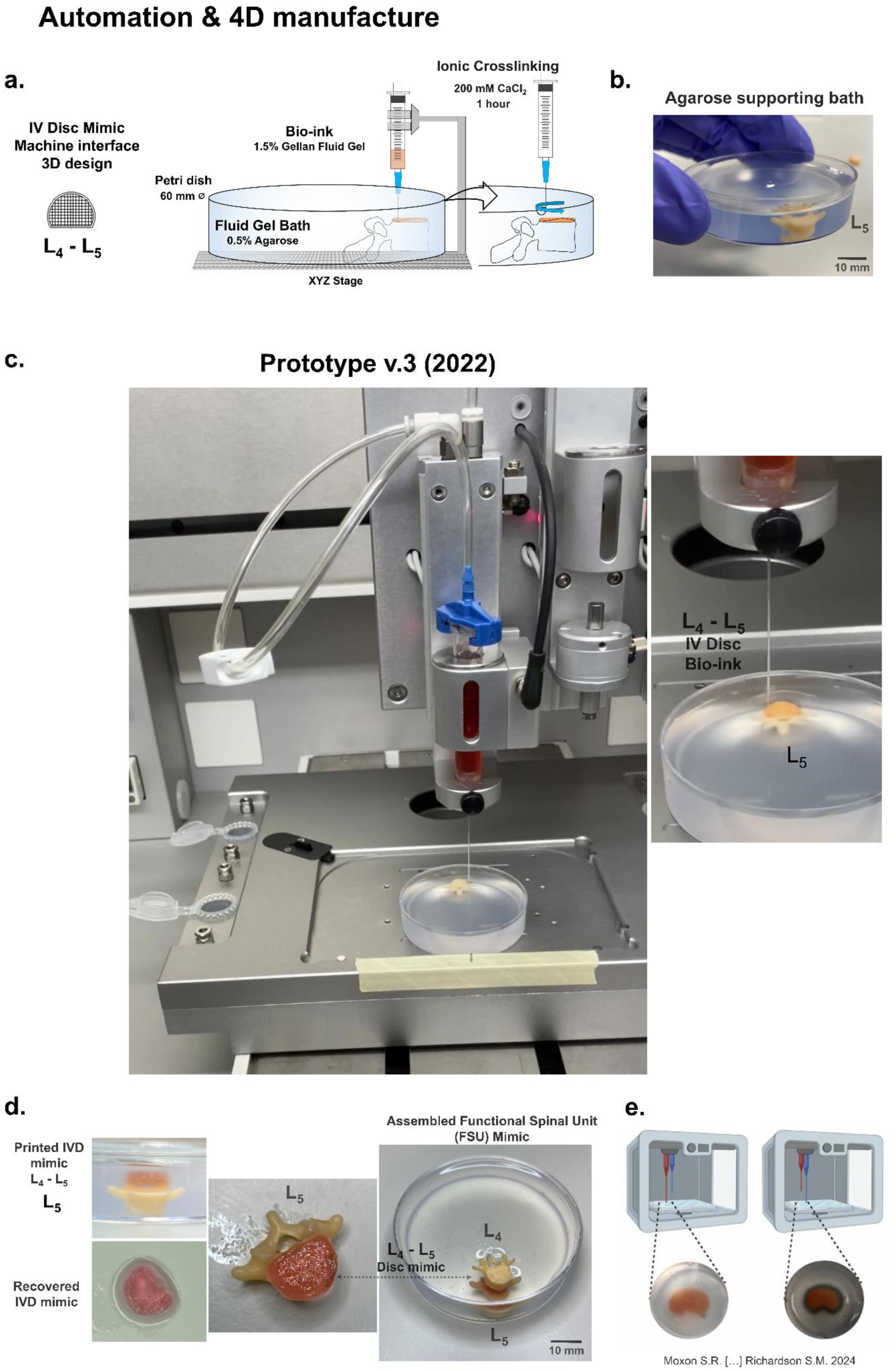
Matrix-supported manufacture - enabling automation and dynamic fabrication. a-b. To prevent gravity-induced deformations, enable verticalisation and improve control of the micro-architecture of the interfaces, we trialled a manufacturing process involving the bioprinting of discs, in conjunction with self-healing, mechanically supporting baths. These provide omnidirectional support throughout the deposition process and enable a cross-linking reaction in situ, during which the bioprinted fluid gel disc (gellan-based) is converted into a structured gel. **c Prototype v.3/2022** enabled 4D functionality through spatially resolved deposition and post-printing maturation and is shown at two stages in the printing process of the L4-L5 interface at the surface of an L5 vertebra, suspended in the matrix for spatial reference. **d** The bioprinted disc structure (**left**, **top**) is successfully aligned with the boundaries of the vertebral body of the L5 vertebra. Following ionic crosslinking induced by injecting a calcium chloride solution around the printed structure, the L4-L5 disc mimic can be recovered from the supporting bath (**left**, **bottom**). The disc can be directly used in the assembly of the spine (**middle**), further processed or re-entered into the printing cycle. The figure on the **right** shows the disc reintroduced as part of a motion segment (L5 vertebra - L5-L4 disc - L4 vertebra) into a supporting bath, to continue the gradual production of a spinal construct (to facilitate the deposition of the L4-L3 disc). **e** Where available, multi-nozzle printers can spatially deliver additional phases of material, as shown in this example from the work of Moxon et al. (2024), which explores the deposition of a collagen-based bioink around a gellan core. Image used under the terms of the CC BY 4.0 license.

#### Matrix-supported manufacture - enabling vertical construction

To prevent gravity-induced deformations and improve control of the micro-architectural complexity of the spinal soft-tissue elements, vertical manufacture of both discs and the spine as a unit could be assisted by automatic methods, such as 3D bioprinting in conjunction with mechanically tuneable support baths (**Figures 10 and 11** **respectively**). These can also enable production at scale and spatial control of cell deposition, if biological material is used. To investigate this, we developed a bioprinting technique assisted by self-healing fluid gel baths to support the disc production process and enable 4D functionality through spatially resolved deposition and post-printing maturation (**Figure 10a-d)**. This manufacturing strategy is also advantageous in a biological fabrication context, when using biotic matter, to preserve viability during the extensive printing times that are typically involved. In this set-up, agarose particulate gel baths (0.5% w/v) loaded into containers of adequate volumes/geometries (**Figure 10b-d**, **Figure 11c-d**), provide omnidirectional support throughout the fabrication process, ensuring that the deposited disc templates, made of low viscosity bioinks, remain stabilised over prolonged printing durations (minutes-hours). This is particularly important in this multi-material system design which requires sequential deposition and curing intervals of bioinks and the addition of vertebral elements.

**Figure 11:**
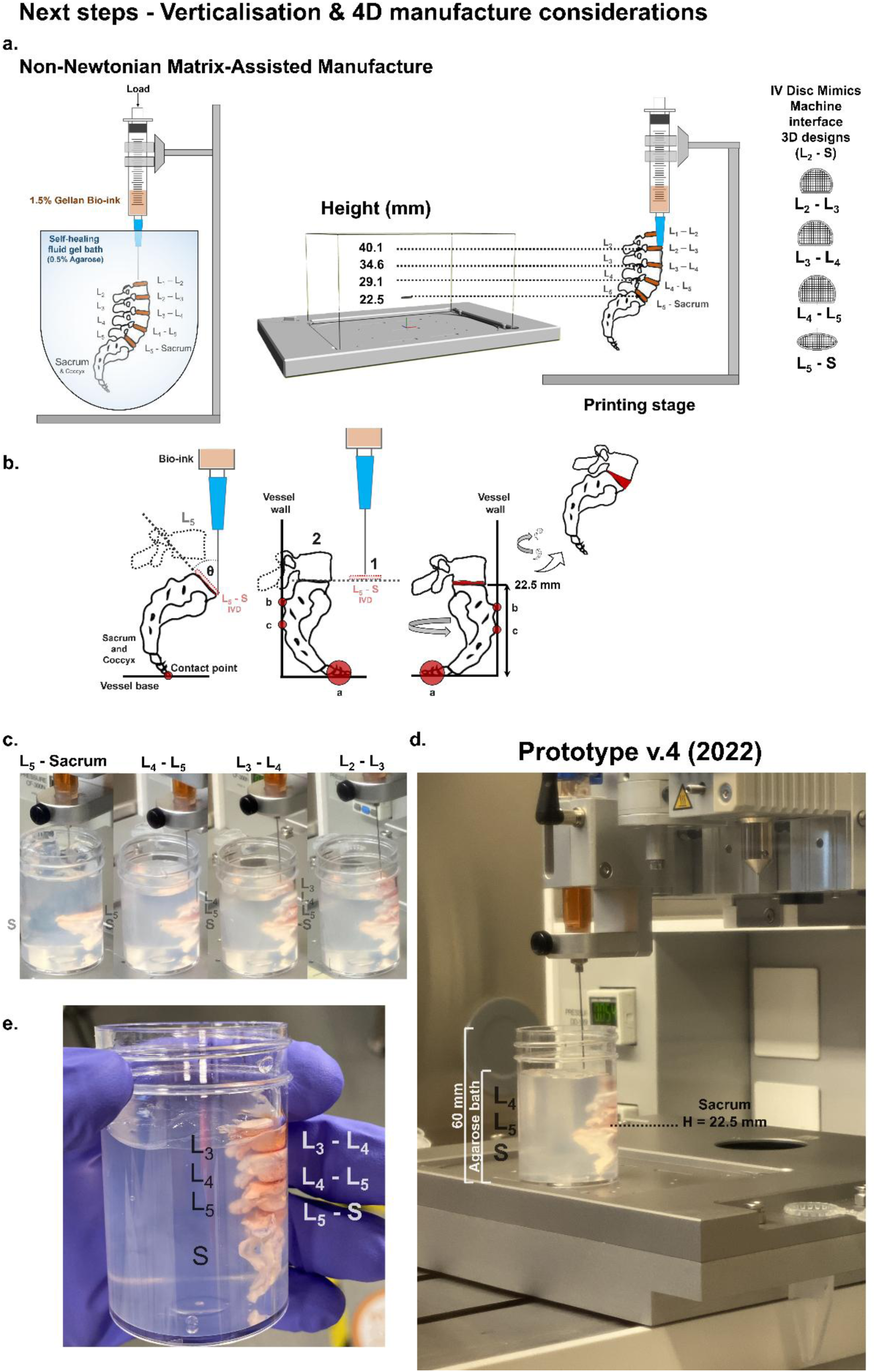
Whole column engineering (3): Verticalisation, spatial mapping and morphogenic fabrication. Spinal manufacture would benefit from a single, unified printing environment which could help embed reactions within a common matrix and provide a shared coordinate system (**a**). A coordinate system for producing a vertical spinal construct from the sacral to the L2-L3 level was developed in this work. **b-c** The sacral/coccygeal bone mimic was the first element introduced in the suspending matrix to establish a stable reference and foundational starting point for the printing process. Subsequently, the L5-S disc interface was printed at the surface of the sacrum, followed by the addition of the L5 vertebra, and the printing of the subsequent interface (L4-L5), continuing up to the L2-L3 interface. This sequential fabrication approach enabled the measurement and refinement of the relevant spatial coordinates/vertical positioning throughout the evolving three-dimensional construct (**Prototype v.4/2022** - **d-e**). **b** To enable printing under the constraints of existing biomanufacturing instruments, the sacrum was manually repositioned into an orientation that allowed planar deposition of the L5-S disc, ensuring compatibility with the printer’s configurations. The new orientation also allowed for the sacrum/coccyx to be supported by the inner wall of the vessel during printing, thereby maintaining stability through increased contact points, without requiring extensive base support. However, this configuration introduced a geometric limitation, due to the extensive processes of the superjacent vertebra (starting with L5), which extend over the sacrum. The structure was therefore rotated 180° to accommodate this constraint and was primarily used as a heigh reference - with subsequent manual readjustment performed post-printing.

Agarose (C_12_H_18_O_9_) is also a bioderived polysaccharide able to produce structured fluids through a similar shear-cooling fabrication technique as described above. It is also a linear polysaccharide, comprised of D-galactose and 3,6-anhydro-L-galactopyranose units [32].

The agarose supporting baths are a suspension of discrete microgel particles dispersed within the continuous phase of this biopolymer. Consequently, these materials exhibit yield-stress behaviour and thixotropy which enables them to self-heal and recover structurally around other phases, such as the low-viscosity 1.5% gellan disc bioinks. Importantly, they can simultaneously allow nozzle movement during bioink deposition while dynamically restructuring and maintaining structural fidelity after printing.

**Figure 10c** presents a manufacturing process involving the biofabrication of individual disc templates as separate structures within their respective support baths, using corresponding vertebrae for guidance and morphometric validation. Disc morphologies were designed in the printing instrument’s BioCAD software (**Figure 10a**, **Figure 11a**), for the lumbar region, focusing on the L_2_-Sacrum/Coccyx area. This methodology is particularly suitable for a gradual assembly of the spinal model and where the disc requires adjustments before addition to the spinal structure.

This prototype (**Prototype v.3/2022 -** **Figure 10c**) is shown at an initial and advanced stage in the printing process, illustrating the successful deposition of the L_4_-L_5_ disc at the surface of an L_5_ vertebra. **Figure 10d** shows an example of the successful alignment of the deposited IVD mimic with the boundaries of the vertebral body of the L_5_ vertebra, which had been placed in the supporting matrix for reference. The disc template was subsequently ionically crosslinked in situ by injecting a calcium chloride solution (200 mM CaCl_2_) around the disc using a fine needle (23 G). Throughout an incubation period (up to one hour), the Ca^2+^ ions in the solution diffused from the injected pockets in the bath through the gellan fluid gel matrix, inducing network formation/gelation by acting as a bridge between negatively charged carboxylate groups (-COO^-^) present on the D-glucuronic acid subunits of neighbouring polymer chains (**Figure 10a**). This process does not chemically affect the supporting bath, as agarose is a neutral polysaccharide which gels primarily through physical mechanisms [33], including hydrogen bonding. The final IVD mimics can be readily retrieved from the bath following the establishment of a cross-linked gel matrix. They can either be processed further as required, or used in the assembly of the spine - in one stage, when all the necessary discs have been produced or through a series of incremental stages. Alternatively, individual motion segments can be re-entered into the printing cycle to enable the gradual assembly and alignment of the spine, as illustrated in **Figure 10d**, which shows an L_5_ vertebra/printed L_5_-L_4_ disc/L_4_ vertebra construct which was re-introduced into a supporting bath to facilitate the deposition of the L_4_-L_3_ disc. This approach would help to preserve hydration and viability of the discs (as well as manoeuvrability of the elements) during the whole-spinal assembly process.

Where available, multi-nozzle printers can spatially deliver additional phases of material, as shown in **Figure 10e**, from the work of Moxon et al. (2024) [15], which explores the deposition of a layer of Type I collagen bioink around a similar concentration gellan disc (1%), and at a relevant scale. These researchers used gellan as the material for the *nucleus pulposus* mimic, (with a 0.5% bovine collagen solution as the *annulus fibrosus*). In contrast, we used this tuneable gel as the scaffold for the overall disc shape/*annulus fibrosus* template. When considered from an alternative manufacturing perspective, printing outer layers of collagen type I around the gellan scaffolds could have potential for further anatomical modelling, also relevant for our own system design. For example, it could be used for building external ligaments (rich in type I collagen) surrounding/stabilising the spine - such as the anterior longitudinal ligament, running along the front and sides of vertebral bodies/discs to provide stability to the spine; as well as the posterior longitudinal ligament running vertically at the back of the vertebral bodies and discs, lining the spinal canal. This would also allow creating an interface and gradient into the *annulus fibrosus* structure, which is externally composed of type I collagen, whereas internally, towards the inner jelly-like disc, becomes more fibrocartilaginous and composed of collagen type II. [34, 35]. More widely, this multi-ligamentous manufacturing approach would also be helpful if printing the spine as a whole unit, as discussed in the next section (and illustrated in **Figure 11**).

Ideally, the vertical manufacture of discs and vertebral integration into a complete spine should be performed within a single, mechanically-supporting bath (**Figure 11a**) rather than using post-printing assembly. The use of a unified printing environment would enable the continuous and highly precise deposition of the disc interfaces within a shared coordinate system, minimising the need for manual intervention and alignment errors. From a chemical perspective, it could help to embed all reactions within a common matrix and provide a more uniform environment and crosslinking conditions. From a manufacturing perspective, this strategy would also enhance efficiency, reduce material waste and minimise processing time. However, this is a much more significant technical challenge compared to printing individual components.

We sought to address this through the development of an initial foundational model that could help us identify the technical challenges, focusing on the lumbar-sacral segment as a starting point (specifically L_2_ - Sacrum/Coccyx). In many respects, this region, combining the mobile spine and the fused sacral bones, and characterised by several anatomical curvatures, is the most difficult to reconstruct experimentally. A range of digital intervertebral disc morphologies was designed as described above, for the L_5_-S, L_4_-L_5_, L_3_-L_4_ and L_2_-L_3_ interfaces. These were designed around the ceramic vertebral correspondents and previous morphometric data and measured, in length and at their longest width: 8.56 mm L x 3.54 mm W (L_5_-S); 7 mm L x 5.8 mm W (L_4_-L_5_); 6.8 mm L x 5.2 mm W (L_3_-L_4_); and 6.1 mm L x 5.5 mm W (L_2_-L_3_).

The first step in the biomanufacturing process was the placement of the sacral/coccygeal bone mimic in the suspending bath to establish a stable reference and foundational starting point for the printing process. Subsequently, the L_5_-S disc interface was printed at the surface of the sacrum, followed by the addition of the L_5_ vertebra, and the printing of the subsequent interface (L_4_-L_5_), continuing up to the L_2_-L_3_ interface (**Figure 11b-c**). This sequential fabrication approach enabled the measurement and refinement of the relevant spatial coordinates/vertical positioning throughout the evolving three-dimensional construct (**Prototype v.4/2022 -** **Figure 11d-e**).

Certain physical modifications had to be made to enable this, due to the curved and inclined geometry of the sacral-coccygeal bone. Firstly, the morphometry of this bone poses challenges for conventional bioprinting instruments (**Figure 11b**). Most commercially available (triaxial) bioprinters (as of 2026) are optimised for planar, layer-by-layer deposition and therefore are not able to print on highly non-planar or angled surfaces. Although recent advances have demonstrated the ability to print on irregular/curved geometries, they are based on bespoke systems developed in-house and are not widely available [36–38]. To enable printing under the constraints of the available systems, the sacrum was manually repositioned into an orientation that allowed planar deposition of the L_5_-S disc, ensuring compatibility with the printer’s configurations. This approach was also advantageous as the sacral structure had limited contact with the vessel at its base, due to the thin and narrow anatomy of the coccyx, which increased the possibility of misalignment during extruder movement, despite the support provided by the bath. The new orientation allowed for the sacrum/coccyx to be firmly supported by positioning it against the inner wall of the vessel during printing, thereby maintaining position and stability through increased contact points, without requiring extensive base support. However, this configuration introduced a new technical challenge, as the superjacent mobile spine elements, starting with L_5_, have large processes which extend over the sacrum and cannot be accommodated beyond the supporting wall. Consequently, the sacrum structure was rotated 180° during printing to accommodate this constraint and was primarily used as a heigh reference - with subsequent manual readjustment performed post-printing. This enabled biofabrication of the segment despite the geometric limitations. However, in future iterations, custom-designed fixtures or sacrificial support structures could be engineered to stabilise the complex lower-sacral geometry, enabling printing to occur more centrally in the vessel and thus accommodate this structure in the relevant conformation. Alternatively, the mobile spine can be fabricated separately, with the sacrum bone mimic applied at the end of the process (with printing commencing at the L_5_-S disc level). Additionally, a strategy for inducing the relevant angular conformation has to be developed during or post-printing. This could be performed in the future either in an automatic manner using advanced -including multi-axial- printers (as they continue to evolve); or semi-automatically, by introducing, post-printing, an external support reinforcement similar to that described above (**Figure 7b**); therefore enabling the correct angular orientations and maintaining spatial positioning of the elements throughout maturation.

A coordinate system for printing the spinal segment was therefore successfully developed through this work and is presented in **Figure 11a**. This work enabled precise spatial mapping and control of print positions within the output area/support bath. This is helpful because it provides guidance for future iterations, delineating precise locations for disc deposition, taking into account the presence of the vertebrae.

#### Assessing the biomechanics of the motion segments

To understand the mechanical behaviour of the individual motion segments comprising the spine prototype (multi-phase disc-vertebral units), we assessed a batch of units from the lumbar section – comprising the L_4_ vertebra /L_4_-L_5_ IVD/ L_5_ vertebra using axial compression testing (**Figure 12**). This allowed assessing the load-bearing response and deformation profile of the unit as a whole. This spinal level was chosen because in the clinical context, it is the most frequently affected segment by degeneration, largely due to its high biomechanical loading [39].

**Figure 12:**
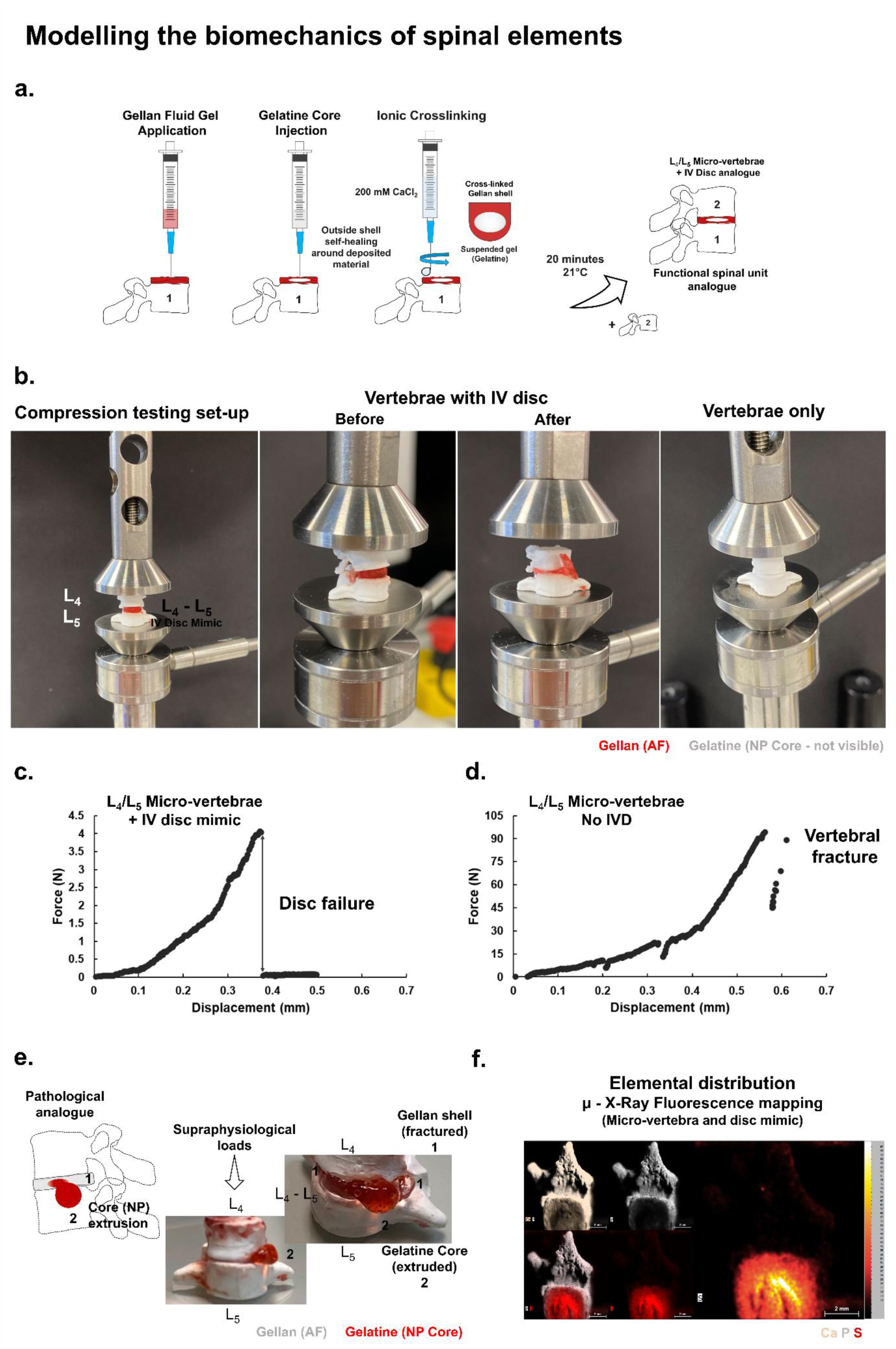
Modelling the tissue composition-structure-function relationship in motion segments. a-b. To understand the mechanical behaviour of the individual motion segments comprising the spinal prototype, we assessed a batch of units from the lumbar section – comprising the L4 vertebra/L4-L5 multi-phase IVD/ L5 vertebra, using axial compression testing. **a** The motion segments were prepared by forming the L4-L5 disc analogues at the surface of a ceramic L5 vertebra, by applying the gellan matrix first, followed by the injection of the gelatine core. Subsequently, the gellan scaffold was cross-linked into a gel by applying a calcium chloride solution in a dropwise fashion, allowing an incubation time of 20 minutes at 21°C. The L4 vertebra was subsequently applied to complete the functional spinal unit analogue. Units without connecting discs (L4 - L5 vertebral constructs only) were also tested for comparison. **b** The compression testing set-up, including the complete motion segment configuration and the vertebrae-only model. **c-d** Two mechanical behaviours were observed in the system. In integrated vertebra-IVD-vertebra constructs, mechanical compression progresses until disc failure is detected in the system (in this example, up to 4.07 N) (**c**). However, when compression is applied to the paired vertebrae model, compression continues to a much greater extent (up to 94.38 N) (**d**), until a terminal fracture is detected in the vertebrae. **e** Calcium-mediated crosslinking enhances the mechanical properties of the *annulus fibrosus* template, which transforms from a gel into a solid mass; but does not chemically affect the gelatine core stored inside. This chemical behaviour is helpful to recreate both the regional distribution of mechanical properties and the pathological behaviour in conditions such as disc prolapse. During supraphysiological loads, the discs in motion segments experience an extrusion of the gelatinous inner core, which pushes through the outer layer through fractures. **f** This protein rich core was also important in the spinal prototype to model a useful tissue biochemistry-structure-function relationship. The Micro-X-Ray fluorescence elemental maps illustrate the spatial distribution and co-localisation of Calcium, Phosphate (as markers for the calcium phosphate mineral content) and Sulphur (as an indicator of organic molecules/protein content) in a multi-phase disc-vertebra construct. Concentrated sulphur deposits can be observed in the *nucleus pulposus* region, with traces present in the *annulus fibrosus* due to the inclusion of the visual tracer (a sulphonated salt of Alizarin), which was also used as a positive control. Throughout the set of tests, the pigment Alizarin Red was used as a label either for the *annulus fibrosus* (**a**-**b,f**) or the *nucleus pulposus* (**e**) to enable visualisation and help validate the mechanical behaviour.

The motion segments were prepared by forming the L_4_-L_5_ disc structures at the surface of a ceramic L_5_ vertebra, by applying the gellan fluid gel matrix first, followed by the injection of the gelatine core, as described above. Subsequently, the gellan scaffold was cross-linked into a gel by applying a calcium chloride solution (60 μl) in a dropwise fashion, allowing an incubation time of 20 minutes at 21°C, in a sealed container. Following this period, the L_4_ vertebra was applied to complete the functional spinal unit analogue (**Figure 12a**). In this set of experiments, the *annulus fibrosus*/gellan template was labelled with a trace of Alizarin Red S pigment to allow observing the morphological changes during testing. Units without connecting discs (L_4_-L_5_ vertebral constructs) were also tested in a stacked conformation to identify differences in mechanical behaviour. **Figure 12b** illustrates the compression testing set-up and a complete motion segment configuration before and after terminal damage.

Two mechanical behaviours can be observed in the system, with examples from each response provided in **Figure 12c-d**. When tested as an integrated vertebral-IVD-vertebral construct, mechanical compression progresses until disc failure is detected in the system (up to 4.07 N) (**12c**). However, when compression is applied to the adjacent vertebrae model, compression continues to a much greater extent (up to 94.38 N) (**12d**), until a terminal fracture is detected in the ceramic vertebrae – also reflecting fractures of the vertebral processes (spinous or transverse), laminae and pedicles, not only the vertebral body.

Although axial compression data for human motion segments is enormously variable, being influenced by demographic factors, pathology, bone mineral content as well as the experimental set-ups (for an analysis, see [40]), for lumbar motion segments, ultimate compressive strengths have been reported in the range of 810 - 10090 N (for males and females in the 17 – 74 years age range), with the L_4_-L_5_ motion segment strengths reported in the range of 1298 N (78 yr female) to 9116 N (24 yr male) [41]. This large variability further highlights the need for personalised bioengineered models for spinal research, which can be mechanically tuned according to the conditions required.

Ca^2+^-mediated ionic crosslinking substantially increases the mechanical properties of the *annulus fibrosus* gellan template, which transforms from a gel into a solid mass; but does not chemically affect the gelatine stored inside, as gelatine’s integrity originates from a physically crosslinked protein network, governed by hydrogen bonding and hydrophobic interactions. This chemical behaviour is helpful to recreate both the regional distribution of mechanical properties in a disc and the pathological behaviour in conditions such as disc prolapse (**Figure 12e**). In this demonstration, axially compressed discs in these motion segments, when exposed to supraphysiological loads, experience an extrusion of the fluid-like cores, which push through the outer layer through fractures.

As described previously, the protein rich core was also important in this prototype to model not only a relevant tissue biochemistry, but also a useful structure-function-biomechanical behaviour relationship (**Figure 12f**).

#### Next-generation vertebral models

With the exception of the vertebral bioceramic model developed here, a tissue engineered model of a vertebra is also missing and represents an area that remains insufficiently addressed. Current vertebral models used in research include anatomical specimens, computational models and synthetic physical models [9, 42, 43]. Several full-scale synthetic models of vertebrae (and the spine) exist [44–49], designed for surgical training, implantology and forensic research, as opposed to tissue engineering, disease modelling and understanding tissue evolution. Current tissue-engineered skeletal models focus on developing mature & osteocyte-containing bone tissue [19, 50], lamellar bone [51] or trabecular bone constructs [52, 53]. Therefore, the osteogenic vertebral model presented here offers a platform for further refinement and complexity. For example, engineering the internal vertebral architecture, mimicking porous trabecular bone, could be useful for modelling the local and regional spinal physiology by enabling cell colonisation, vascularisation and -in motion segments- nutritional transfer to the intervertebral space.

Maturing technologies such as the 3D printing of ceramics (extrusion or photopolymerisation-based) represent one method which could facilitate the incorporation of complex internal architectures, such as the trabecular-like lattice. This internal structure can be easily digitally customised, as shown in **Figure 13** using an L_5_ vertebra, using standardised, relevant, infill patterns, such as a 3D honeycomb structure (in this example, with an infill density of 15% as a starting point).

**Figure 13:**
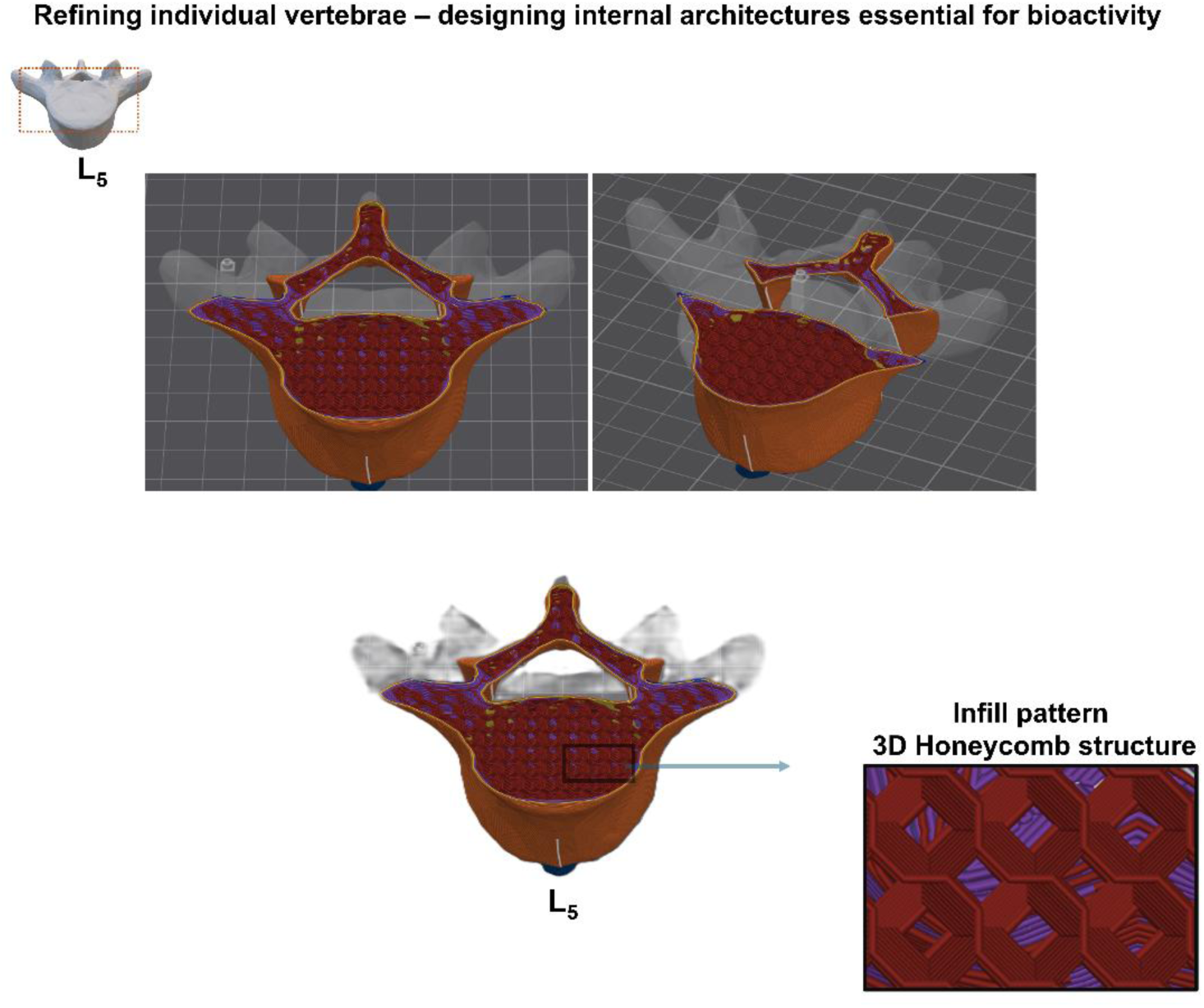
Enhancing the vertebral architectures. Vertebrae could be converted into functional metabolic units through structural modifications of the internal architecture, which would enable cellular infiltration and creating an environment permissive for angiogenesis. This could be enabled by maturing technologies, such as ceramic 3D printing. These modifications would be useful not only for designing an active vertebral unit/a tissue engineered model of a vertebra, but also for supporting the viability of surrounding discs, which in the physiological context rely on the vertebral supply of nutrients. A starting point would be digitally introducing trabecular-like architectures using easily customisable pre-defined lattices available in 3D manufacturing software – in this case, a relevant 3D honeycomb lattice, exemplified using cross-sections through an L5 vertebra.

The ceramic vertebrae can therefore be turned into metabolically active elements through structural modifications (openings and cavities) and exploiting the osteoconductive properties of the material, which could promote cellular infiltration, angiogenesis and diffusion of nutrients to adjacent structures (endplates and disc mimics).

Furthermore, for biomedical applications, the ceramic vertebral manufacturing process could be adapted to produce vertebrae with clinically relevant mechanical properties, focused on long-term functional stability in a corrosive physiological environment, using more bioinert ceramics (e.g. hydroxyapatite - Ca_10_(PO_4_)_6_(OH)_2_, alumina - Al_2_O_3_, zirconia - ZrO_2_ or other technical ceramics [54, 55]). Achieving this would require leveraging new and evolving forms of ceramic additive manufacturing. **Figure 14a** presents an example of an in-house developed ceramic 3D printing instrument based on direct ink writing (DIW) additive manufacturing, which we used for printing the first cervical vertebra (C_1_/Atlas), using a zirconium silicate base (ZrSiO_4_) and a carboxymethyl cellulose (CMC) binder system. The vertebra was printed at the anatomical scale using a 1 mm nozzle. This technology is ideal for producing relevant internal architectures as discussed above, mimicking the trabecular bone porosity (simulated here using a 50% grid structure infill), and at the true scale (**Prototype v.5/2026**). A current limitation of extrusion-based ceramic printing methods is the reduced ability to produce miniaturised structures due to the rheological properties of ceramic pastes and slurries, which require relatively large nozzle diameters for deposition. This is because, in order to achieve sufficient green-body density and ensure shape fidelity following deposition, there is a need for high solid loading (in this case, 85.6 wt%), which requires high pressure to sustain flow through the small nozzle. These factors can limit the minimum feature size that can be printed.

**Figure 14:**
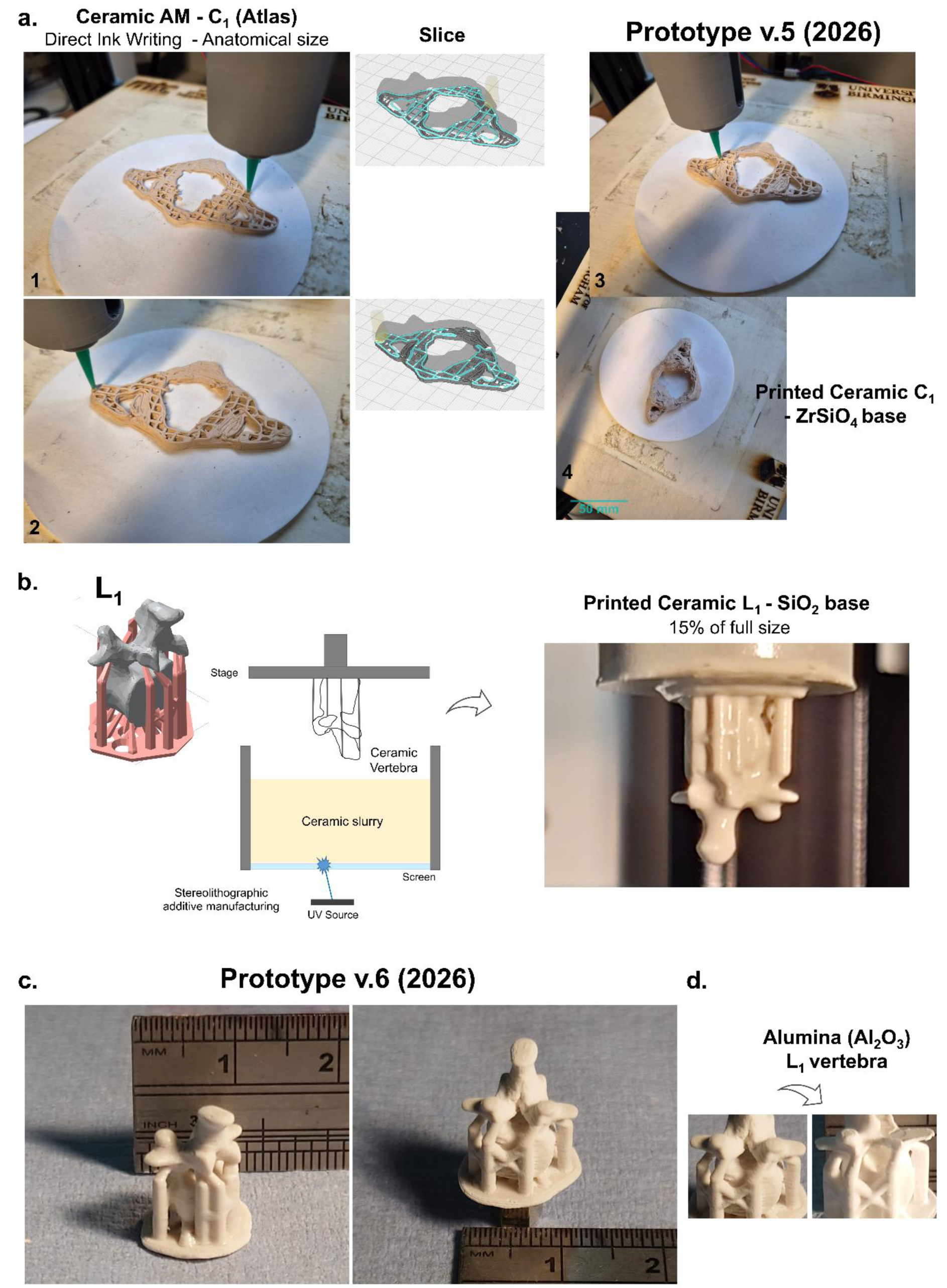
Fabrication of advanced vertebrae using emerging and maturing ceramic printing technologies. **a** An internally developed system for Direct Ink Writing additive manufacturing of ceramics was employed to print the first cervical vertebra (Atlas/C1) with a simulated porous internal architecture, at the anatomical scale, using a zirconium silicate base (**Prototype v.5/2026**). The vertebra is shown at intermediate and final stages in the printing process (**1**-**4**). **b-c** An internally developed system for stereolithographic additive manufacturing of ceramics was used to 3D print the first lumbar vertebra (L1), using a silica-based ceramic mixture, at the required scale (15%). The vertebra was fabricated using a modified instrument similar to that employed in the first part of the work, enabling the direct production of the ceramic vertebra, and with an excellent surface finish. This manufacturing process removes the requirement for an intermediate polymer-printing step followed by ceramic casting (**Prototype v.6/2026**). **d** We are currently optimising this process for application with the common medical implant material Alumina (Al2O3).

Among evolving ceramic additive manufacturing technologies (which can also address this limitation), stereolithographic (photochemical) ceramic 3D printing is one of the most promising types due to its ability to produce complex, scaled-down geometries with good surface quality and precision. **Figure 14b-c** presents an example of a lumbar ceramic vertebra (L_1_) that we produced using an in-house built system, at the required scale (15%), using a silica-based ceramic slurry (SiO_2_) in a blend of acrylate monomers, dispersant and photoinitiator. The L_1_ vertebra was fabricated using a modified instrument similar to that described in the first part of the work (**Figure 4c**, **Figure 5a**), enabling the direct fabrication of the ceramic vertebra and removing the requirement for an intermediate polymer-printing step followed by ceramic casting (**Prototype v.6/2026**). We are currently optimising this process for application with the common implant material Alumina (Al_2_O_3_)(**Figure 14d**), widely used in orthopaedic medical implants [56] and some interbody cages for spinal fusion surgery [57, 58].

#### Conclusions and outlook

The spine is a kinetic and physiological system, where mechanical strain is not uniform, tissue maturation can vary in different regions and signalling is spatially distributed.

Bioengineering models reflecting these complexities is essential for biomedical research, reducing the reliance on animal testing and exploring pathological processes, as well as the refinement of medical interventions.

Developing models at the reduced, in vitro scale (e.g. 15%) can help produce biotic models, provide controlled and scalable experimental conditions and overcome several technical limitations associated with studying the full spinal organ (and in vivo). Technical advantages include precise control over environmental conditions, accessibility for real-time observation and continuous monitoring as well as higher throughput testing ability, particularly important for mechanistic studies.

This is also an exceptionally complex engineering challenge, which requires the convergence of advanced biomaterials, active matter and sophisticated biomanufacturing strategies.

The work presented here looked at the core engineering questions that can define how a vertebral column-system can be built, stabilised and made to function as an integrated physiological system (and under gravitational load). It focused on anthropometric, biophysical, biochemical and manufacturing optimisations that can generate a spinal system – an axial, multi-segment, multi-phase biomechanical construct. The core prototype includes geometrically accurate mineralised vertebral units assembled in sequence into an anatomical, columnar architecture and mechanically connected by bioactive interfaces.

Through several variations of this prototype (which collectively span a ten-year development period), each benefitting from emerging technological innovations, we assessed whether spinal tissue analogues can maintain structure and function when organised as a continuous, bioactive and mechanically connected system. Considerations also included system-level adaptations, longitudinal mechanobiology and multi-segment coupling, which are phenomena that cannot be studied in single disc or functional spinal unit models. The prototype therefore enables transitioning from local to distributed mechanics, static to evolving properties, isolated to interacting tissues and controlled outputs to adaptive behaviour.

The model introduces, for the first time, an axial, spinal architecture, where spatial organisation, tissue maturation and load transmission are allowed to co-evolve. Importantly, it enables studying the interaction between living matter and spinal organisation; as well as coupled vertebral and disc tissue maturation across segments. This is particularly useful for exploring disease mechanisms in conditions such as Scoliosis or Ankylosing Spondylitis, where deformities are associated with simultaneous biological and structural changes within both vertebrae and discs. The prototype can help to assess how evolving mechanical asymmetries (which are not designed into the system) can emerge from bioactivity or tissue growth, mechanical feedback and structural (re)organisation.

The model could also be further enriched with key cell-types, including pathological phenotypes and *nucleus pulposus* and *annulus fibrosus* derived cells.

In developmental models, the prototype could help draw conclusions about the cellular dynamics, how mineralisation in one region can affect neighbouring regions, whether these mechanical changes propagate along the column and how biological maturation becomes spatially patterned.

As shown in these tests, the model is complex enough to show the emergence of system-level behaviour. Even in a simplified, baseline model, the constructs can simulate non-linear behaviour, load redistribution between segments and region-specific mineralisation patterns, which are essential for simulating both developmental and pathological states.

These design features inevitably introduce additional behaviours such as local stiffening effects and emerging mechanical gradients as well as coupled adaptation between motion segments, which could be assessed in future experiments.

Future work could focus on enhancing tissue and anatomical complexity (e.g. vertebral internal architecture, mechanically optimising the discs), personalising the model and integrating additional anatomical structures - such as the anterior and posterior longitudinal ligament mimics discussed above. Further technical work could include assessing the adaptive architecture to identify whether alignment stabilises over time, the structure self-reinforces under load or, more widely, how it self-organises under mechanical constraints. Another focus area is the intervertebral space – improving integration between discs and vertebrae and studying interface evolution when biologically active, mechanically active or both.

These models could also help to validate assumptions used in computational spine models, acting as a biological validation system for simulation work.

## Materials and Methods

### Manufacturing high-resolution polymer vertebrae

The 3D digital files (.obj files) for individual vertebrae were acquired from the Database Center for Life Science (Japan) under the CC-BY-SA 2.1 licence (http://lifesciencedb.jp/bp3d/) (now widely available from additional online sources). Files were processed and prepared for stereolithographic UV printing in the ELEGOO Mars Instrument’s accompanying ChiTuBox slicing software (version 1.5.0).

The 7 cervical (C_1_-C_7_), 12 thoracic (T_1_-T_12_), 5 lumbar (L_1_-L_5_) vertebrae, as well as the sacrum and coccyx mimics, were digitally scaled-down for printing to 15% of their actual size. Each of the vertebrae was sequentially arranged on the virtual stage, supported by lower-density pylon-shaped scaffolds, to ensure the attachment of both vertebral bodies and processes to the motorised stage during printing. The support size was set to ‘medium’, with a contact diameter of 0.8 mm and depth of 0.4 mm, respectively. The connection shape was selected as conical and the density of the pylons was set to 50%. To produce high-resolution physical vertebrae, the 3D codes were loaded into an ELEGOO Mars UV Photocuring LCD 3D printer (Shenzhen, China), which produced the solid geometries from a methacrylate resin that was cured, layer by layer, when exposed to a UV source (wavelength of 405 nm), as the samples were progressively lifted from the resin tank. The metal tank contained a transparent 140 x 200 mm, 0.15 mm thick FEP replaceable film lining the bottom, offering a 95% light transmittance. Following printing, the vertebrae were retrieved from the stage by carefully detaching from the pylons and gently washed with 95% isopropanol to remove any uncured resin. All the procedures were conducted inside a chemical fume hood.

### Duplication of the vertebral and whole-spine anatomical morphologies

Two commercially available elastomeric impression materials were used for producing copies of the cervical, thoracic, lumbar vertebra and the sacrum-coccyx. The first type consisted of a liquid silicone base and its curing agent (Beckly, UK). To optimise and identify the ideal formulation for duplication and recovery of the anatomical structures, different ratios of the compound to its curing agent were prepared to determine which variation enabled the best resolution, whilst offering sufficient flexibility to allow recovery of the encased vertebra without damaging the delicate transverse processes. A ratio of 1.5:0.5 curing agent to silicone elastomer by mass was found to produce optimal properties and was used throughout the work. 12 grams of the liquid silicone mixture was poured over each vertebra, placed at the centre of a diamond-shaped weighing boat (7 cm d_1_ x 5 cm d_2_). The batch of vertebral moulds was allowed to set for 24 hours, following which the individual vertebrae were carefully extracted through small surface openings created by the vertebral projections.

A second impression material (a commercially available, putty-type elastomer) was used to replicate flat and complex vertebrae (first cervical vertebrae, L_5_ and the sacrum/coccyx), which were more difficult to duplicate/recover. The formulation consisted of a silicone moulding paste and its curing agent (Siligum, Pebeo, France). A 1:1 ratio by mass was used, which generated high-definition imprints with a rapid setting time (approximately 5 minutes).

This material was also used to duplicate the assembled spine structure. The macroscopic template was applied to a layer of elastomer of 1.5 cm height, weighing 45 grams.

### Production of the bioceramic replicates

The cavities generated by the polymeric templates within the impression materials were subsequently filled with a biomimetic calcium phosphate cement paste to generate biochemically relevant mineral vertebrae. A ratio of 2 g of a fine β-TCP powder (Ca_3_(PO_4_)_2_; <125 μm particle size) to 1 ml of orthophosphoric acid (3.5 M H_3_PO_4_) was used to generate the cement paste. The liquid mixture was poured into the cavities of the moulds, on a vibrating platform, to encourage uniform filling of the shapes. All reagents were cooled to 4°C before the commencement of the work to slow down the setting time of the cement, which forms rapidly due to the exothermic reaction between the calcium phosphate powder and orthophosphoric acid.

### Production of the disc interfaces

Gellan bioinks (1.5% w/v) were used as surrogates for the *annulus fibrosus*. A commercial grade of low acyl gellan powder (Kelcogel^®^, Arcinova, UK) was dissolved in deionised water to a concentration of 1.5% w/v whilst simultaneously being heated up to 120°C and agitated at 700-800 rpm using a 7 cm rod-shaped magnet, on a magnetic stirrer hot plate, to encourage dissolution. A 500 ml volume of the material was prepared in a 1 litre capacity borosilicate glass reagent bottle (Fisherbrand^TM^, Thermo Fisher Scientific, USA) designed for elevated-temperature applications. Once dissolved, the liquid became optically clear and the heating was switched off. The solution was subsequently shear-cooled to room temperature for up to 6 hours. During testing, 0.1g of Alizarin red S pigment (Alfa Aesar, UK) was added to the gel to provide a visual contrast.

To simulate the inner compartment of vertebral discs, the *nucleus pulposus* was modelled using gelatine (food-grade, Oetker Group, Germany). 9.25g (±0.5g) of suine-derived gelatine sheets were re-hydrated in deionised water for 10 minutes in preparation for dissolution. The sheets were subsequently re-suspended into 250 ml of deionised water and were gently agitated, while the container was submerged in a water bath set to 37°C to assist with a slow, uniform dissolution.

In the original (pathological) prototype (**v.1/2017**), osteogenic interfaces were produced by generating a reaction in situ between the physiological plasma components fibrinogen and thrombin, leading to the formation of a fibrin gel. The reactions were produced inside six-well microplates (34.8 mm well diameter), which had been pre-coated with 1.5 ml of the colourless silicone elastomer Sylgard184 (elastomer base: curing agent ratio of 10:1). Bovine thrombin (200 U/ml in F12K media - *see below*) was used as part of a DMEM culture medium solution (containing 10% FBS), at a ratio of 50 μl/ml, alongside the antifibrinolytic agents aminohexanoic acid (200 mM stock concentration) and aprotinin (10 mg/ml stock concentration) at a ratio of 2 μl/ml. Bovine fibrinogen (≥75% of clottable protein - Sigma-Aldrich, USA), prepared in F12K Nutrient Mixture (1×) with Kaighn’s Modification (Gibco, Life Technologies, USA), was added at a ratio of 200 μl (20 mg/ml stock concentration). Gels were allowed to polymerise for 30 minutes during incubation at 37°C, inside a 5% CO_2_ humidity incubator. Bone marrow mesenchymal stromal cells (murine), suspended in culture media, were added at a ratio of 100.000 cells per vertebral-disc construct.

### Bioprinting of interfaces

Disc interfaces were bioprinted within supporting agarose baths (0.5% w/v) using a RegenHu 3DDiscovery system (Villaz-Saint-Pierre, Switzerland), composed of an integrated bioprinting station within a biosafety cabinet enclosure. Discs were printed using a nozzle/needle diameter of 0.39 mm, an operating pressure of 0.07 MPa, a feed rate of 30 mm/s, layer height of 0.31 mm and strand width of 0.45 mm, corresponding to a volumetric deposition rate Q ≈ 4.19 mm^3^/s and an average material velocity inside the nozzle v ≈35 mm/s. Disc geometries were designed in the BioCAD software. Spatial coordinate generation, printing regimes and 3D visualisation was undertaken in the BioCAM software/slicer.

### Non-Newtonian supporting baths

Agarose (C_12_H_18_O_9_) (genetic analysis grade, >1200 g/cm^2^ gel strength) was used for making the non-Newtonian gel baths. For preparing a fluid gel bed, a 0.5% w/v agarose solution (Fisher Scientific Bioreagents, USA) was developed as described above, in dH_2_O by heating to 120°C with magnetic stirring (using a 7 cm rod-shaped stirrer) until dissolution, followed by cooling to 21°C under a constant shear (approximately 700 rpm) for up to 6 hours. The gel may also be produced via an alternative process suitable for use with living matter, which similarly involves shear cooling the solution, following autoclaving through a 121°C cycle (saturated steam, for 15-20 minutes, under pressure of at least 1.1 Bar G). Fluid gels were loaded into either 60 mm diameter petri dishes or 60 mm height/ 60 ml reagent containers and allowed to set for 15 minutes prior to the start of the experiment. The spinal bone mimics were inserted in the correct spatial position using metallic tweezers.

### X-Ray Analysis of the lumbar-sacral spine

X-Rays of a detached lumbar-sacral spine were acquired using a micro-computed tomography system, to image the 3D internal organisation of this multi-phase construct at the micron-scale resolution. A micro-computed tomography system (SkyScan 1172, Bruker Instruments, Germany), was used to scan the spinal section, placed inside a 60 mm diameter cell culture dish, on a rotating stage inside the machine, located at a distance of 260.65 mm from the source. A Hamamatsu X-Ray source was used to scan the sample, operating at a voltage of 80 kV and a current of 100 μA. 2D cross-section slices of the spinal section were acquired using an 11 Mp X-ray camera with a 9.01 μm pixel size, generating images of 27.05 μm pixel size. X-Ray images were captured with a rotation step of 0.2°, with 360° rotation, two frames averaging per step and an exposure time of 1000 ms.

### Micro-X Ray Fluorescence analysis

A micro X-ray fluorescence system (M4 Tornado, Bruker Nano Gmbh, Berlin, Germany) was used to generate spatially resolved elemental maps of discs and vertebrae, using the localization of Ca, P, and S in constructs. The machine contains a rhodium μ-focus X-ray tube and a polycapillary lens, used to focus the X-rays to a spot size of 25 μm. Recordings were taken without sample processing, at room temperature and under vacuum conditions. The X-ray tube voltage used was 50 kV and tube current was 400 μA. For the vertebral-disc constructs, X-Ray fluorescence maps were acquired using a 20 μm spot distance, and 5 ms per pixel exposure time. For the assessments of vertebral batches, X-Ray fluorescence maps of L_4_ and L_5_ vertebrae were acquired using a 20 μm spot distance, and 3 ms per pixel exposure time. Elemental maps were formed in real time by integrating the photon counts around the emission lines of: calcium (Kα1 3.692 keV), phosphorus (Kα1 2.010 keV), sulphur (Kα1 2.309 keV), generating an image where pixel intensity was proportional to the number of X-ray counts/second per electronvolt (eV) from each measured point on the construct. Thus, pixel intensity increased with X-ray counts, with maximum pixel intensity normalized to the highest count rate per eV for each element of interest, across the entire sample.

### Mechanical characterisation of motion segments

Motion segments composed of the L_4_ vertebra/L_4_-L_5_ IVD/ L_5_ vertebra were prepared by injecting 20 μl of a gelatine biopolymer into 50 μl of a suspending gellan polymer at the surface of an L_5_ vertebra. Subsequenltly, the gellan scaffold was cross-linked into a gel by applying a 200 mM CaCl_2_ solution in dH_2_O (60 μl) in a dropwise fashion, allowing an incubation time of 20 minutes at 21°C, in a sealed container. This was followed by the application of the L_4_ vertebra to complete the unit. The segments, as well a variation containing only the vertebral elements for comparison, were subjected to axial compression using a Bose ElectroForce^®^ 5500 instrument (Massachusetts, USA). A preload of 0.2 N was applied. Compression was applied at a rate of 0.005 mm/sec, to a preset level of 1 mm, using a 200N load cell. Control for the instrument and testing regime set-ups were enabled by a WinTest^®^ Digital Control System. Data was captured using the WinTest^®^ 7 software. Compression data was plotted as positive values. A minimum of three motion segments were tested in each group.

### Printing of ceramic vertebrae

#### Direct Ink Writing additive manufacturing of ceramic vertebrae

Ceramic paste feedstocks were prepared from zirconium silicate powder, an aqueous carboxymethyl cellulose (CMC) binder system and magnesium acetate as a minor additive. A 5 wt% CMC binder concentration was selected for the printing trials, with the final paste formulation consisting of 85.6 wt% zircon, 14.0 wt% binder and 0.4 wt% magnesium acetate. Zircon was used as a representative technical ceramic model system. The 85.6 wt% solids loading was selected to provide a high ceramic content while ensuring extrudability and green body shape retention for direct ink writing (DIW) additive manufacturing.

Printing was carried out using a modified Creality CR-20 Pro platform adapted for syringe-based DIW of ceramic pastes. The original polymer extrusion system was replaced with a custom syringe printhead, in which a NEMA 23 stepper motor and lead screw applied controlled displacement to the syringe plunger. Motion and extrusion were controlled using a Duet 3 6HC control board running RepRap Firmware, allowing room temperature deposition of the aqueous ceramic paste without a heated nozzle or build chamber.

The paste was deposited through a 1 mm nozzle using a 0.5 mm layer height and 1.0 mm line width. The geometry was sliced using 50% grid structure infill.

#### Stereolithographic additive manufacturing of ceramic vertebra

A silica-based ceramic slurry in a blend of acrylate monomers, dispersant and photoinitiator was prepared in a dual axis centrifugal mixer (Algimax II GX300, Monitex Industrial Co.) for 120 seconds. The Voxeldance Tango slicer software and the “Bar support-Figure” script using default settings was used to generate the support structure and 3D printing program. The component was printed on a modified ELEGOO Mars 4 DLP vat photo-polymerisation resin printer using a 50 µm layer thickness and 8 s exposure time. Following printing, the part was cleaned in ultrasonicated ethanol for 10 seconds at room temperature. An early alumina-based (Al_2_O_3_) prototype was produced for comparison by adapting an established printing method [59]. Calcined alumina powder (ultra-fine,1.0 µm particle size) (Logitech, Glasgow, UK) was mixed at a solids loading of 76.54% with a monomer blend consisting of equal amounts of HDDA & TMPTA together with dispersant (Disperbyk-111) and Photoinitiator (BAPO) added at 2% and 1% respectively by weight of the total slurry.

## Acknowledgements

The principal investigator (Dr Alexandra Iordachescu) and co-authors would like to thank the National Centre for the Replacement Refinement and Reduction of Animals in Research (NC3Rs – UKRI, United Kingdom) for supporting this research through the award of two research grants to Dr Iordachescu (Grants no. NC/S001859/1 and NC/X000907/1).

The authors would also like to thank Oliver Barella and Ezzat Ahmadouk for supporting the experimental work during the initial phase of the project, as part of an undergraduate research project, designed and supervised by Dr Iordachescu (2019/2020). We would also like to thank Dr Lucy Arkinstall for technical support with mechanical testing.

The skeletal specimen image in Figure 5f is an original photograph of a specimen displayed at the Natural History Museum, London, originally used in the PI’s doctoral thesis (2018). We would like to thank the NHM for allowing photography of their collections.

## Author contributions

Dr Alexandra Iordachescu conceived the work, acquired the funding, designed and performed the research, collected and analysed the data & interpreted the results. Dr Aleksander Cendrowicz and Renush Vigneswaran provided experimental support and expertise with the ceramic additive manufacturing work (design and implementation). Professor Anthony D. Metcalfe and Professor Liam M. Grover contributed with scientific expertise. Dr Aleksandar Atanasov provided technical support and expertise with the biomanufacturing work. The manuscript was written and edited by Dr Iordachescu with input from all authors.

